# Immunologic insights into the critical epitopes of HIV-1 and structure-based characterization of cross-reactive antibodies

**DOI:** 10.1101/2025.07.15.663531

**Authors:** Deepika Jaiswal, Sheenam Verma, Zaid K Madni, Gagandeep Kaur, Kanwal J Kaur, Dinakar M. Salunke

## Abstract

HIV-1 escape from neutralizing antibodies even in the presence of strong host immunity is associated with variations in envelope proteins that drive antigenic diversification. The virus exploits the error-prone nature of reverse transcriptase and the high mutation rate as key survival strategies. However, the rate of production of new variations occurs at relatively slow pace. Interestingly, the immune system often produces cross-reactive antibodies, with anticipated role in neutralizing point mutations in HIV surface proteins by cross-reacting with mutants and tolerating them. In light of this paradox, we explored the mystery of immune evasion and antibody promiscuity by screening single chain variable fragment (scFvs) antibodies against several crucial HIV-1 gp41 epitopes using a phage display library. These findings underscore the broader significance of cross-reactive antibodies. Here, high-affinity cross-reactive scFvs showed physiologically relevant affinities with peptide epitopes, their analogs, and the native HIV-1 gp41 protein. We determined the crystal structure of a high-affinity cross-reactive scFv, DE94, and obtained insights into the molecular interactions of scFv antibodies with peptide epitopes and its natural mutants using molecular docking studies. This analysis of cross-reactive antibodies could contribute to the therapeutic development against immune-evading pathogens and paves the way for innovative strategies for combating viral infections, including emerging global threats.

## Introduction

The immune system produces antibodies to defend the body against invading pathogens by neutralizing them. Traditionally, antigen-antibody interactions were as perceived as highly specific [1]. However, accumulating structural data has revealed that host-pathogen interactions are much more complex and exist in dynamic equilibrium. While the immune system is under pressure to eradicate pathogens, a selection pressure exists on the pathogens to evade detection and establish infection [2]. Fast-mutating viruses such as HIV-1 (Human Immunodeficiency Virus) and influenza exploit this balance by incorporating genetic variations on their surface-exposed envelope proteins and evade the immune response. As a result, the immune system can no longer detect mutated antigenic surfaces, disturbing the equilibrium between the host and the pathogen. The immune system must employ some strategy to defend against these viruses and maintain this equilibrium. Several studies have shown that in the case of infection, the regular B-cell repertoire consists of two types of antibody populations: monospecific and multispecific [3]. While the role of monospecific antibodies is well known, as antibodies are specifically generated against invading pathogens. However, the significance and function of multispecific antibodies remain less understood and require further investigation.

Multispecificity in the affinity-matured antibodies has been previously reported [4], suggesting a potential role for these antibodies in combating the effect of antigenic variations in fast-mutating viruses. Such viruses present a physiologically relevant model to understand the role of multispecificity in the immune response. A previous study on the influenza hemagglutinin (HA) protein and its mutants investigated the mechanisms of antibody cross-reactivity [5], suggesting that the multispecific antibodies may help prevent immune evasion despite some mutations successfully driving antigenic variation. In the present study, we sought to determine if the function of multispecific antibodies was limited to the influenza virus or if similar mechanisms occur in response to other viral infections. To gain a deeper understanding of the functions of multispecific antibodies, we extended our investigation to examine multispecific antibodies targeting HIV-1.

The HIV envelope proteins play a central role in viral replication and are key determinants of the virus’s antigenic and pathogenic properties [6–8]. The functional envelope is a trimer of two heterodimeric surface glycoproteins - gp120 and gp41. gp120 is essential for the attachment of the virus, while gp41 mediates the fusion of the viral and the host membranes. The host’s key defense mechanism to combat HIV infection is antibody-mediated neutralization and many broadly neutralizing antibodies have been identified and studied so far [9, 10]. It has been observed that the gp41 region of the envelope is more conserved than the gp120 [11–13]. Within the gp41 region, one of the elusive vaccine targets is the membrane proximal external region (MPER), outside the viral membrane [14]. Apart from this, the Kennedy epitope is another immunogenic site within the cytoplasmic tail (CTT) of gp41 inside the viral membrane which makes it less accessible. However, numerous studies have demonstrated that antibodies directed against the epitope found in the CTT region can neutralize the virion [15, 16] indicating that specific epitopes become accessible during viral conformational changes. Such antibodies can block the cell-to-cell transmission of HIV by blocking fusion [17–19].

The contrast between HIV immune evasion strategies and recent studies on antibody promiscuity is fascinating. HIV escapes immune surveillance by incorporating mutations in its envelope proteins. It has been reported that even a point mutation is enough to evade the immune response [20, 21]. However, multispecific antibodies are also found to be increased during viral infections [22], revealing an interesting dichotomy. In light of this, our study aims to address the mechanisms underlying the HIV-1 immune evasion and antibody promiscuity. We focused on two critical epitopes within gp41 to understand the role of multispecific antibodies in different hotspots of HIV. Using human-based antibody phage display libraries, we screened antibodies against these epitopes and successfully isolated a diverse panel of cross-reactive antibodies against both epitopes. The selected antibodies showed diversity in their binding affinity, ranging from micromolar to sub nanomolar affinity. Intriguingly, antibodies screened against peptide epitopes were able to recognize the epitope within full-length protein. Crystallographic and *in silico* analysis have provided insights into the molecular mechanism of the interaction of antibodies with the peptides and their variants. Our findings emphasize the physiological relevance of antibody promiscuity in the natural immune response and its potential implications for therapeutic and vaccine strategies targeting HIV.

## Results

### Selection and design of epitopes

The HIV-1 envelope protein comprises of two heterodimers, both of which have been considered the targets for generating monoclonal antibodies. The gp41 is particularly important for vaccine development as it plays a crucial role in membrane fusion. During the transition from pre-fusion to post-fusion state, gp41 undergoes conformational changes, transiently exposing the previously buried epitopes [23–25]. Structurally, gp41 consists of an N-terminal region fusion peptide (FP), followed by N-terminal and C-terminal heptad repeats (NHR and CHR), membrane proximal external region (MPER), transmembrane (TM) and a cytoplasmic tail (CTT) [26] (Figure. 1A). To understand the effect of antigenic diversity on the immune response, we selected two key neutralizing epitopes within gp41. By analyzing the envelope protein in the HIV database genome browser, we selected a 7-mer neutralizing epitope ^738^GERDRDR^744^ in CTT and a 6-mer neutralizing epitope ^662^ELDKWA^66^ in MPER, both recognized for their neutralization potential [15, 16] (Figure 1A). The GERDRDR sequence is a part of the Kennedy epitope (^724^PRGPDRPEGIEEEG**GERDRDR**S^745^), which has been implicated in virus neutralization, incorporation of the envelope in the virion and viral infectivity. Although no broad neutralizing antibodies have been identified against this epitope, antibodies targeting this region can block cell-to-cell transmission by blocking the fusion [17–19]. In contrast, ELDKWA is the core epitope of 2F5, a well-known broadly neutralizing antibody against HIV [27]. Among the several broadly neutralizing antibodies against the MPER region, such as 10E8, LN01, 2F5, and 4E10, the 2F5 antibody binds to the linear N-terminal MPER epitope that had more natural mutants in comparison to other antibodies [11, 28]. The other MPER-directed antibodies recognize helical epitopes located in the C-terminal of the MPER region near the TM region [29].

**Figure 1.**
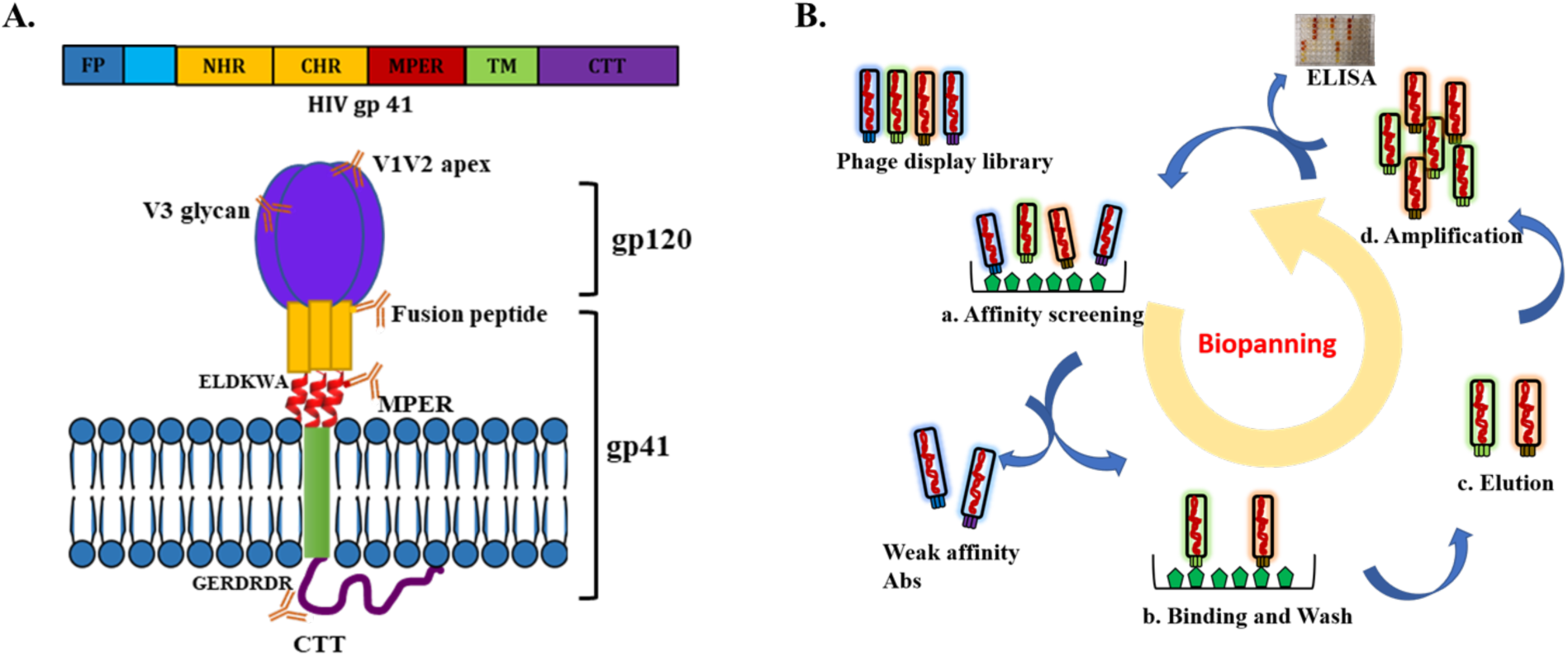
Epitope selection and antibody biopanning led to epitope-specific phage clones. **A.** Schematic illustration of the gp41 domains consisting of fusion peptide (FP), N-heptad repeat (NHR), C-heptad repeat (CHR), membrane-proximal external region (MPER), transmembrane domain (TM), and cytoplasmic tail (CTT). Broadly neutralizing antibodies and their corresponding epitopes are depicted on the gp120 and gp41 proteins of the HIV envelope. **B.** Schematic of phage display antibody library screening, consisting 4 main steps: (a) Affinity screening involves immobilization of BSA conjugated antigen on immunotubes and addition of antibody libraries; (b) High-affinity antibodies bind to the antigens while the weak binders washed off using PBST; (c) Bound phages are eluted for the recovery of high binders followed by (d) Amplification of the eluted epitope-specific phages. Amplified phages from each round are used as input phages for the next round of biopanning.

To understand the extent of degeneracy in antibodies raised against epitopes GERDRDR and ELDKWA, we selected variants from the HIV sequence database using the web based AnalyzeAlign tool (www.hiv.lanl.gov). Based on parameters such as percentage abundance, the impact of mutations, and sequence coverage, we selected six variants for each epitope. To represent sequence variability comprehensively, we included single, double, and triple mutants and each mutation was selected such that it changed the amino acid hypdropathy (Table 1). To identify the epitope-specific antibodies, we focused on using a human phage display library to screen selected epitopes (Figure 1B).

**Table 1.** Selection of neutralizing epitope and its natural escape variants from NIH HIV database.

| S. No. | Mutations | Sequence | % Abundance |  |  |
| --- | --- | --- | --- | --- | --- |
|  |  |  | Group B | Group C | Group A1 |
| 1 | Native epitope | GERDRDR | 37.17 | 0.29 | 0.91 |
| 2 | Single mutant | RERDRDR | 0.02 | - | - |
| 3 |  | GGRDRDR | 0.98 | - | 1.82 |
| 4 |  | GEQDRDR | 5.21 | 41.64 | 16.82 |
| 5 | Double mutant | GEQDRGR | 1.91 | 4.51 | 11.82 |
| 6 | Triple mutant | GEQGRGR | 0.21 | 0.44 | 10 |
| 7 |  | GGRDSGR | 0.1 | - | - |
| 1 | Native epitope | ELDKWA | 57.52 | - | 6.98 |
| 2 | Single mutant | ALDKWA | 11.52 | 0.3 | 75.24 |
| 3 |  | ELDKWD | 2.62 | 0.32 | - |
| 4 |  | EWDKWA | 0.49 | - | - |
| 5 | Double mutant | KLDEWA | 1.36 | - | - |
| 6 |  | ALDKWQ | - | 0.32 | 0.47 |
| 7 | Triple mutant | ALGKWD | 0.06 | 28.73 | - |

### Antibody biopanning and analysis of cross-reactivity for GERDRDR epitope Isolation and characterization of GERDRDR-specific antibodies

Antibodies can target various regions of the HIV envelope protein involved in virus infection and entry. The GERDRDR epitope is involved in membrane fusion during viral entry. Four rounds of biopanning were performed to isolate epitope-specific antibodies using the human-based semi-synthetic Tomlinson I and J scFv phage display libraries. Progressive enrichment of epitope-specific phages was observed in rounds 3 and 4 (Table 2) as confirmed by polyclonal phage ELISA (Figure S1). A total of 400 monoclones from the fourth round of biopanning were tested for binding to the native epitope and a varying degree of binding reactivity was observed (Figure 2A). Based on peptide/BSA OD490 values, 100 top binders were selected and further analyzed by ELISA (Figure S2A).

**Figure 2.**
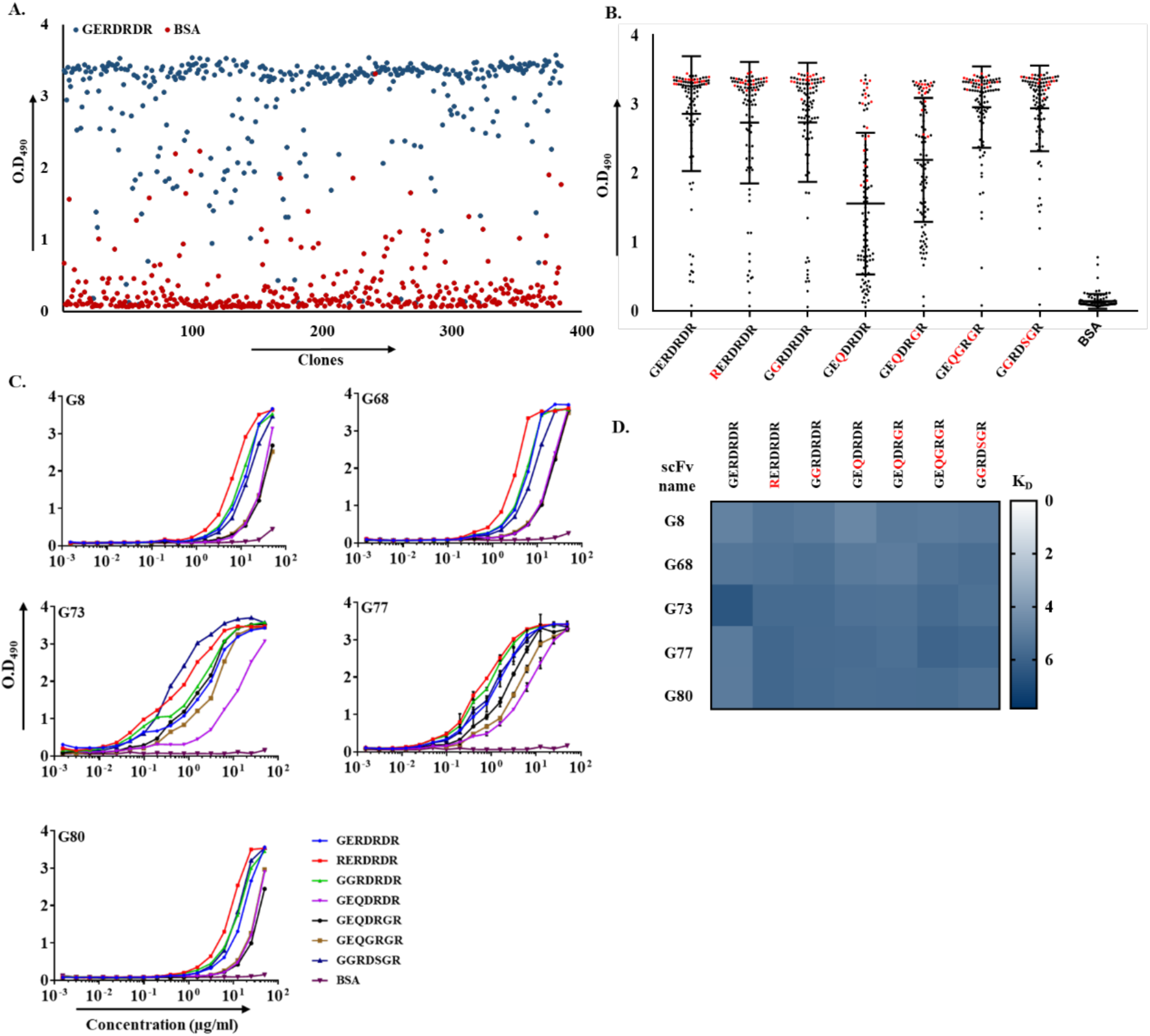
Binding analysis of monoclonal phages against GERDRDR and their cross-reactivity. **A.** Identification and isolation of GERDRDR-specific monoclonal phages by ELISA. **B.** Cross-reactivity profile of the selected phage clones against all the mutants **C.** Each scFvs’ binding capacity to GERDRDR and its variants was tested in indirect ELISA, showing a dose-response curve. Binding graphs are combined per antibody. **D.** Heat map displaying the binding of scFvs to the natural escape variants of the GERDRDR epitope. The -log10 KD values range for each epitope is indicated from 0 to 7, and BSA was used as a negative control.

**Table 2.**
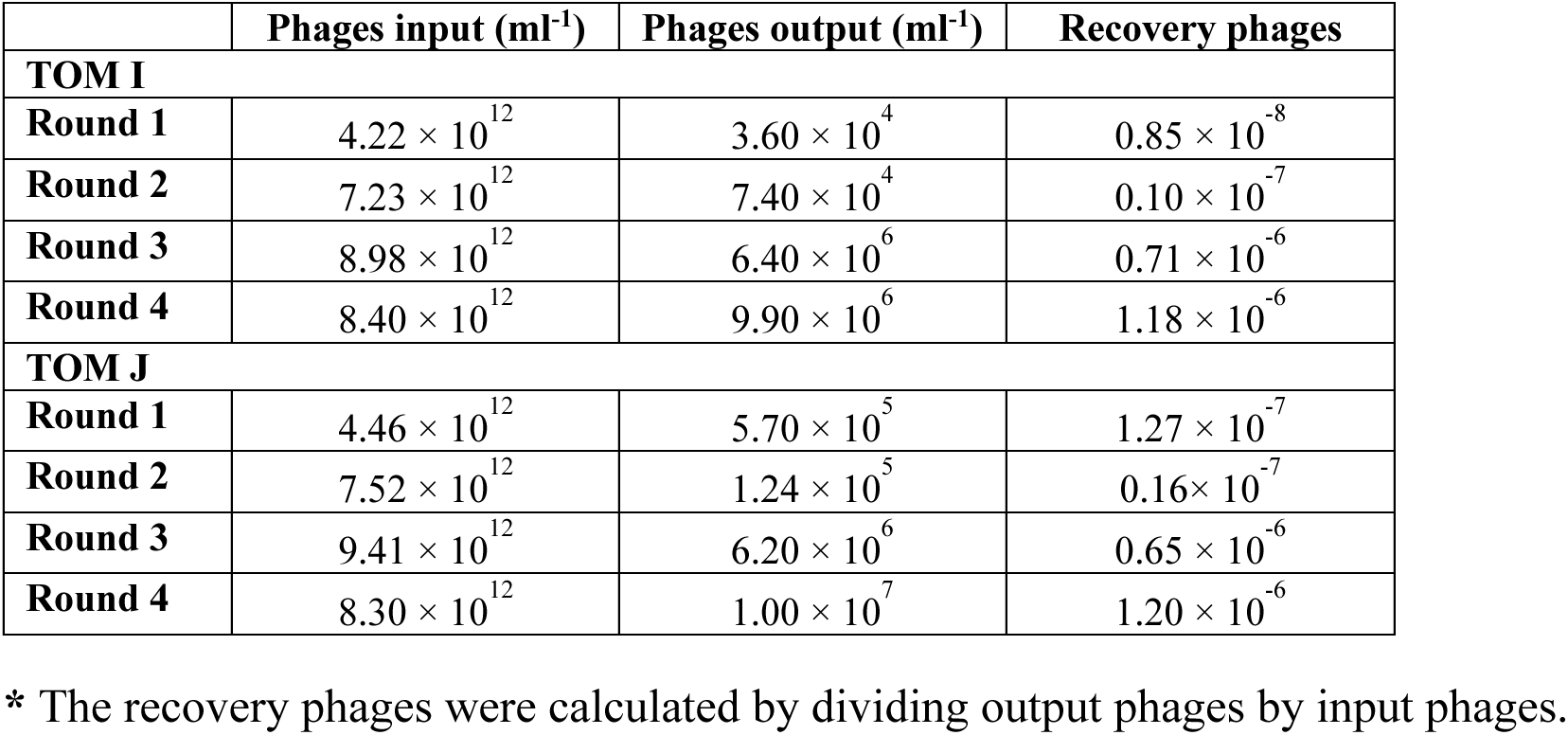
Enrichment of epitope-specific output phages in the successive rounds of screening.

To assess the extent of cross-reactivity, monoclonal phages with high binding to native epitope were tested against the cross-clade variants. These phages were examined for their ability to cross-react with single mutants (RERDRDR, GGRDRDR, GEQDRDR), double mutants (GEQDRGR), and triple mutants (GEQGRGR, GGRDSGR) (Figure 2B). While most mutants showed no significant change in mean OD values compared to the native epitope, except for GEQDRDR and GEQDRGR, which showed reduced binding. (Figure. S2B). This reduced binding suggests that specific amino acid substitutions at the 3rd and 6th positions of the epitope can significantly impact the interaction with the antibodies. Sequencing of 15 cross-reactive high binding clones resulted in the identification of five unique scFv (Figure S2C & S3). These scFvs were cloned into the pET22b vector, expressed, and purified to high homogeneity (Figure S4A and B).

To assess the binding of these selected scFvs to both the native and variant epitopes in soluble form, ELISA was performed in a series of dilutions ranging from 50 µg mL^-1^ to 1.52 ng mL^-1^. All five scFvs showed binding to full panel of variants (Figure 2C), although with different binding range. EC50 values of scFv G8, G68, G73, G77, and G80 were determined to be 11.72 µM, 5.51 µM, 2.07 µM, 1.26 µM, and 16.60 µM respectively. Among all the scFvs, G77 showed the highest affinity followed by G73. Further analysis using Bio Layer Interferometry (BLI) confirmed the multi-reactive potential and binding kinetics of scFvs to HIV epitope mutants (Table 3). A heat map of the equilibrium dissociation constant (KD) showed that all five scFvs bind in the micromolar range (Figure. 2D) reinforcing their potential as broadly reactive antibodies capable of recognizing epitope variants associated with HIV immune evasion.

**Table 3.** BLI data showing KD values of all the scFvs against GERDRDR, its variants, and gp41 protein.

|  | GERDRDR | RERDRDR | GGRDRDR | GEQDRDR | GEQDRGR | GEQGRGR | GGRDSGR | gp41 |
| --- | --- | --- | --- | --- | --- | --- | --- | --- |
| **G8** | $16.74 \times 10^{-6}$ | $5.74 \times 10^{-6}$ | $8.16 \times 10^{-6}$ | $25.77 \times 10^{-6}$ | $7.69 \times 10^{-6}$ | $9.19 \times 10^{-6}$ | $7.19 \times 10^{-6}$ | $6.86 \times 10^{-6}$ |
| **G68** | $5.97 \times 10^{-6}$ | $4.32 \times 10^{-6}$ | $3.37 \times 10^{-6}$ | $8.26 \times 10^{-6}$ | $9.99 \times 10^{-6}$ | $4.08 \times 10^{-6}$ | $2.92 \times 10^{-6}$ | $2.85 \times 10^{-6}$ |
| **G73** | $3.44 \times 10^{-7}$ | $2.20 \times 10^{-6}$ | $2.20 \times 10^{-6}$ | $3.91 \times 10^{-6}$ | $4.21 \times 10^{-6}$ | $3.68 \times 10^{-6}$ | $3.66 \times 10^{-6}$ | $5.21 \times 10^{-6}$ |
| **G77** | $8.98 \times 10^{-6}$ | $1.54 \times 10^{-6}$ | $2.28 \times 10^{-6}$ | $2.51 \times 10^{-6}$ | $3.42 \times 10^{-6}$ | $1.22 \times 10^{-6}$ | $1.67 \times 10^{-6}$ | $2.83 \times 10^{-6}$ |
| **G80** | $9.40 \times 10^{-6}$ | $1.53 \times 10^{-6}$ | $2.18 \times 10^{-6}$ | $2.86 \times 10^{-6}$ | $2.68 \times 10^{-6}$ | $2.56 \times 10^{-6}$ | $3.73 \times 10^{-6}$ | $3.50 \times 10^{-6}$ |

### Binding analysis of GERDRDR-specific antibodies to gp41 and gp160

It was evident from the affinity data that scFvs could bind the native peptide epitope and its variants in a physiologically relevant range. To assess the binding in a more native context, we next evaluated the binding of scFvs with recombinantly expressed full-length gp41containing the native epitope sequence. Biotinylated gp41 protein was immobilized on the streptavidin (SA) sensor, and different scFvs were used as the analyte at concentrations starting from 10µM to 70 nM. Interestingly, all the scFvs exhibited binding to gp41 protein in the micromolar range (Table 3). The binding affinity curves with association and dissociation profiles are shown in Figure 3A.

**Figure 3.**
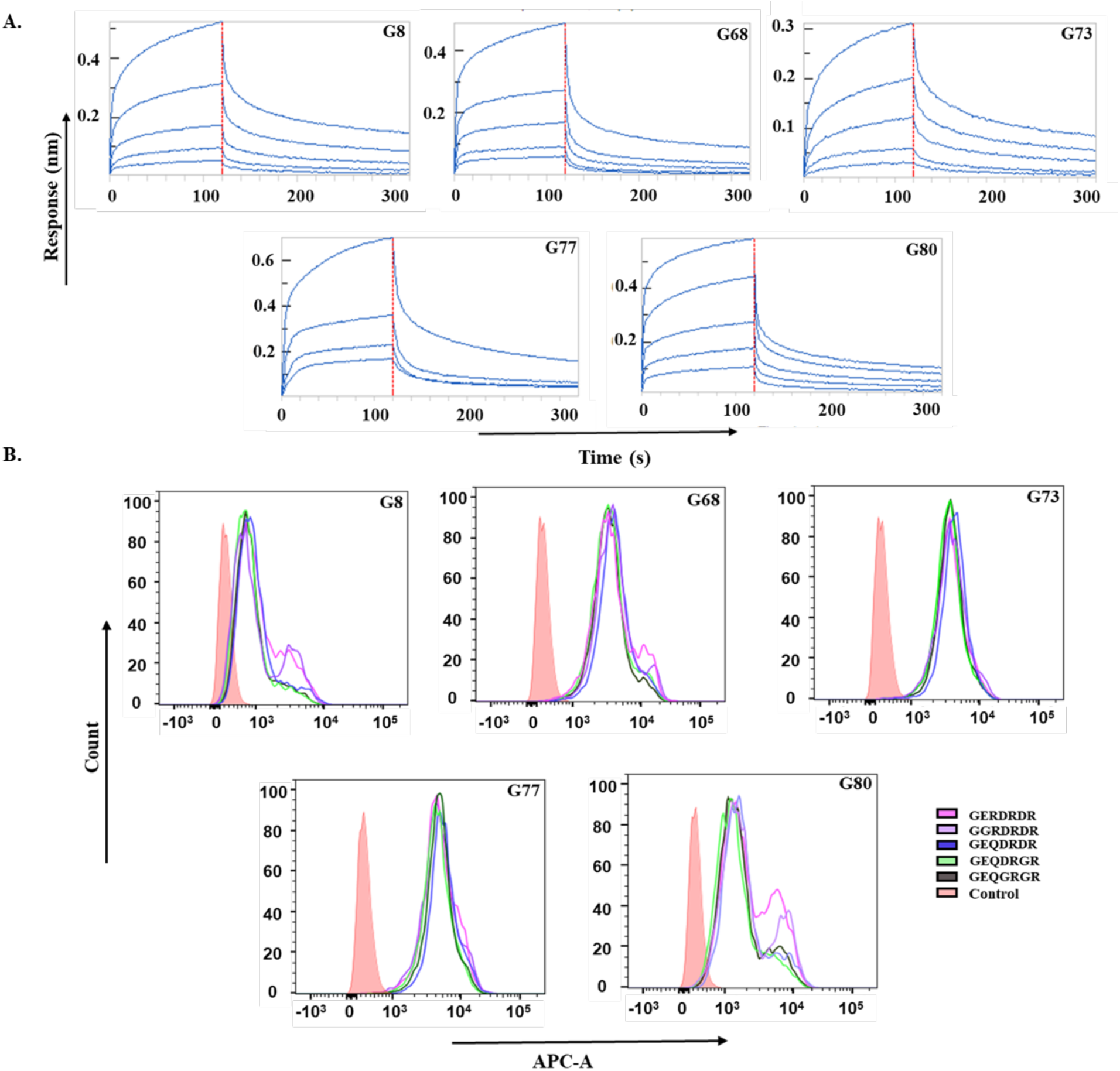
Binding analysis of scFvs to the gp41 and gp160 proteins. **A.** Sensogram showing binding of gp41 protein (ligand) immobilized on the SA sensor to the scFvs (analyte) at different concentrations. The sensogram shows the association and dissociation profiles of all the scFvs. **B.** All the scFvs were analyzed for their binding with the gp160 protein expressed on HEK 293T cells by flow cytometry. Unstained cells were used as a control.

To further investigate whether the scFvs could recognize the native conformation of gp160 in a membrane-anchored state, we tested the binding of the scFvs to the full-length gp160 protein expressed on the mammalian cell surface. The expression of the gp160 protein on the cell surface will approximately correspond to a model of the virus particle in which gp160 is embedded in the viral membrane. Site-directed mutagenesis was performed to make variants of full-length gp160 protein using HIV SF162 plasmid. The two variants (RERDRDR and GGRDSGR) could not be obtained by site-directed mutagenesis due to technical challenges. Flow cytometry analysis revealed that all the scFvs were able to recognize the native gp160 protein and four of its variants as evident by the shift in the fluorescence peak (Figure 3B). The shift in the whole peak with reference to the control peak indicates strong binding to the gp160 protein. The higher the shift in peak, the stronger the binding. The disparity in the peak shift with different scFvs indicated different binding strengths. Overall, these results indicated that the scFvs can recognize both recombinant and membrane bound forms, highlighting their multi reactive potential.

### Antibody biopanning and analysis of cross-reactivity for ELDKWA epitope Isolation and characterization of ELDKWA-specific antibodies

The membrane fusion associated ELDKWA epitope, exposed on the exterior of the viral membrane, was also subjected to screening against Tomlinson I, Tomlinson J, and HuScL4 libraries as previously discussed (Figure 1B). Phages specific to this epitope were enriched for 4-5 rounds of biopanning (Table 4), with increased binding was observed after each round by polyclonal ELISA (Figure. S5). From the fourth and fifth rounds, 1150 monoclonal phages were tested for binding to the native epitope, and 150 high binders selected based on the peptide/BSA OD490 value (Figure 4A, S6A). Of these, 80 top binders were selected for further analysis.

**Figure 4.**
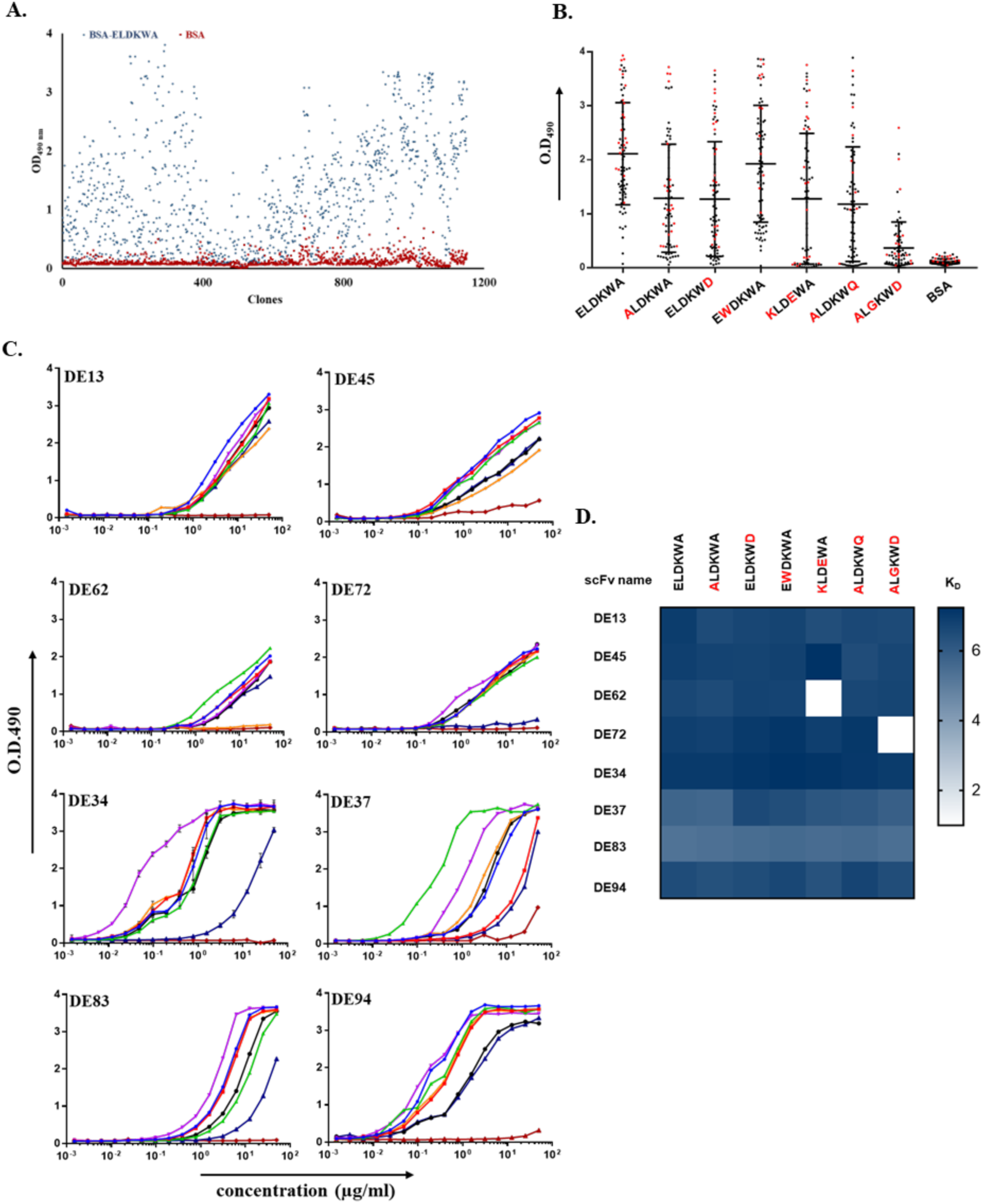
Binding analysis of selected monoclonal phages exhibiting cross-reactivity. **A.** Identification and isolation of epitope-specific monoclonal phages by ELISA. **B.** Cross-reactive profile of the selected phage clones against all the mutants. **C.** Each scFvs’ binding capacity to ELDKWA and its variants was tested in an indirect ELISA. Binding graphs are combined per antibody. **D.** Heat map representing the binding affinities of scFvs to ELDKWA epitope and its natural escape variants. BSA was used as a negative control. Binding strength is represented by - log10 KD values in the range of 0-7.

**Table 4.** Table showing enrichment of epitope-specific output phages in successive rounds of screening.

|  | **Input Phages**<br>**($\text{mL}^{-1}$)** | **Output Phages**<br>**($\text{mL}^{-1}$)** | **Recovery phages** |
| --- | --- | --- | --- |
| **TOM I** |  |  |  |
| **Round 1** | $2.61 \times 10^{12}$ | $8.00 \times 10^5$ | $3.06 \times 10^{-7}$ |
| **Round 2** | $3.60 \times 10^{12}$ | $1.60 \times 10^6$ | $4.50 \times 10^{-7}$ |
| **Round 3** | $2.02 \times 10^{12}$ | $3.62 \times 10^8$ | $1.79 \times 10^{-4}$ |
| **Round 4** | $2.82 \times 10^{12}$ | $8.41 \times 10^8$ | $2.98 \times 10^{-4}$ |
| **TOM J** |  |  |  |
| **Round 1** | $2.44 \times 10^{12}$ | $4.96 \times 10^6$ | $2.03 \times 10^{-6}$ |
| **Round 2** | $4.77 \times 10^{12}$ | $1.48 \times 10^7$ | $3.10 \times 10^{-6}$ |
| **Round 3** | $3.01 \times 10^{12}$ | $1.92 \times 10^9$ | $6.40 \times 10^{-4}$ |
| **Round 4** | $2.78 \times 10^{12}$ | $1.34 \times 10^9$ | $4.80 \times 10^{-4}$ |
| **HuScL4** |  |  |  |
| **Round 1** | $2.44 \times 10^{12}$ | $1.60 \times 10^5$ | $6.55 \times 10^{-8}$ |
| **Round 2** | $8.80 \times 10^{12}$ | $1.28 \times 10^6$ | $1.45 \times 10^{-7}$ |
| **Round 3** | $3.60 \times 10^{12}$ | $3.20 \times 10^7$ | $8.8 \times 10^{-6}$ |
| **Round 4** | $1.20 \times 10^{12}$ | $2.5 \times 10^8$ | $2.08 \times 10^{-4}$ |
| **Round 5** | $5.70 \times 10^{12}$ | $2.50 \times 10^9$ | $4.38 \times 10^{-4}$ |

To comprehend the breadth of antigen recognition, the selected 80 phages were tested for binding to six selected variants. Remarkably, most phage monoclones recognized majority of variants demonstrating strong cross-reactivity (Figure 4B). In comparison to the native epitope defined as percentage O.D. value of 100%, triple mutant ALGKWD showed the lowest mean value of 13%, single mutant EWDKWA exhibited a mean value of 92% that was comparable to the natural epitope. Single mutants, ALDKWA and ELDKWD, showed a mean OD490 value of 59%, while that of double mutants, KLDEWA and ALDKWQ, had 59% and 54%, respectively (Figure S6B). These results indicate that phages screened against the native peptide epitope were highly cross-reactive with the escape variants. Sequencing of 20 high binders yielded four distinct clones from the HuScL4 library and four from the Tomlinson libraries (Figure S6C & S7). These scFvs were cloned into the pET22b vector, expressed, and purified (Figure S8A, B).

The binding affinities were assessed by ELISA across various dilutions, ranging from 50.00 µg mL^-1^ to 1.52 ng mL^-1^. Interestingly, the selected eight scFvs showed binding to the native epitope with varying affinities. The EC50 values for scFvs DE13, DE45, DE62, DE72, DE34, DE37, DE83, and DE94 ranged from 4.27 µM to 0.25 µM, indicating a spectrum of binding affinities spanning from micromolar to submicromolar -range. Notably, scFvs DE94 and DE34 exhibit the highest binding affinity among all scFvs (Figure 4C). While scFvs DE13, DE34, DE37, DE83, and DE94 recognized all - tested variants, scFv DE62 failed to bind KLDEWA and scFv DE72 did not recognize ALGKWD (Figure 4C) indicating the differential binding specificities among the clones.

To complement the affinity calculations by ELISA, BLI was used to quantify the binding kinetics. BLI analysis revealed the diversity in binding affinities among the scFvs (Figure 4D, Table 5). Some scFvs retained the binding across all the selected variants while others lost binding to double or triple mutants indicating the sensitivity to the sequence variations.

**Table 5.** Comparative analysis of KD values of all the scFvs against native epitope and its variants.

| scFv | ELDKWA | ALDKWA | ELDKWD | EWDKWA | KLDEWA | ALDKWQ | ALGKWD | gp41 |
| --- | --- | --- | --- | --- | --- | --- | --- | --- |
| DE13 | $2.17 \times 10^{-7}$ | $3.13 \times 10^{-7}$ | $2.35 \times 10^{-7}$ | $1.95 \times 10^{-7}$ | $3.98 \times 10^{-7}$ | $2.66 \times 10^{-7}$ | $2.99 \times 10^{-7}$ | $5.08 \times 10^{-6}$ |
| DE45 | $1.38 \times 10^{-7}$ | $1.85 \times 10^{-7}$ | $1.92 \times 10^{-7}$ | $1.59 \times 10^{-7}$ | $0.58 \times 10^{-7}$ | $3.52 \times 10^{-7}$ | $2.31 \times 10^{-7}$ | $6.46 \times 10^{-6}$ |
| DE62 | $2.49 \times 10^{-7}$ | $3.20 \times 10^{-7}$ | $1.80 \times 10^{-7}$ | $1.98 \times 10^{-7}$ | No binding | $2.42 \times 10^{-7}$ | $2.10 \times 10^{-7}$ | $1.55 \times 10^{-5}$ |
| DE72 | $1.39 \times 10^{-7}$ | $1.62 \times 10^{-7}$ | $0.96 \times 10^{-7}$ | $0.76 \times 10^{-7}$ | $1.39 \times 10^{-7}$ | $0.82 \times 10^{-7}$ | No binding | $3.39 \times 10^{-6}$ |
| DE34 | $11.90 \times 10^{-8}$ | $9.70 \times 10^{-8}$ | $7.43 \times 10^{-8}$ | $5.76 \times 10^{-8}$ | $6.84 \times 10^{-8}$ | $8.71 \times 10^{-8}$ | $1.09 \times 10^{-7}$ | $2.76 \times 10^{-6}$ |
| DE37 | $1.95 \times 10^{-6}$ | $2.27 \times 10^{-6}$ | $2.85 \times 10^{-7}$ | $4.40 \times 10^{-7}$ | $7.20 \times 10^{-7}$ | $8.35 \times 10^{-7}$ | $1.36 \times 10^{-6}$ | $2.67 \times 10^{-6}$ |
| DE83 | $5.51 \times 10^{-6}$ | $4.49 \times 10^{-6}$ | $3.44 \times 10^{-6}$ | $2.81 \times 10^{-6}$ | $3.41 \times 10^{-6}$ | $2.95 \times 10^{-6}$ | $3.99 \times 10^{-6}$ | $3.64 \times 10^{-6}$ |
| DE94 | $3.44 \times 10^{-7}$ | $4.78 \times 10^{-7}$ | $3.68 \times 10^{-7}$ | $2.48 \times 10^{-7}$ | $5.25 \times 10^{-7}$ | $2.00 \times 10^{-7}$ | $3.73 \times 10^{-7}$ | $0.94 \times 10^{-6}$ |

### Binding analysis of ELDKWA-specific antibodies to gp41 and gp160

As described previously, it was necessary to check whether the scFvs screened against the peptide epitope could also recognize the corresponding full-length HIV protein. Binding kinetics with gp41 protein confirmed that all scFvs were capable of binding to gp41 with KD values in the micromolar range (Figure 5A, Table 4). The highest binding affinity observed was 0.94 µM for scFv DE94.

**Figure 5.**
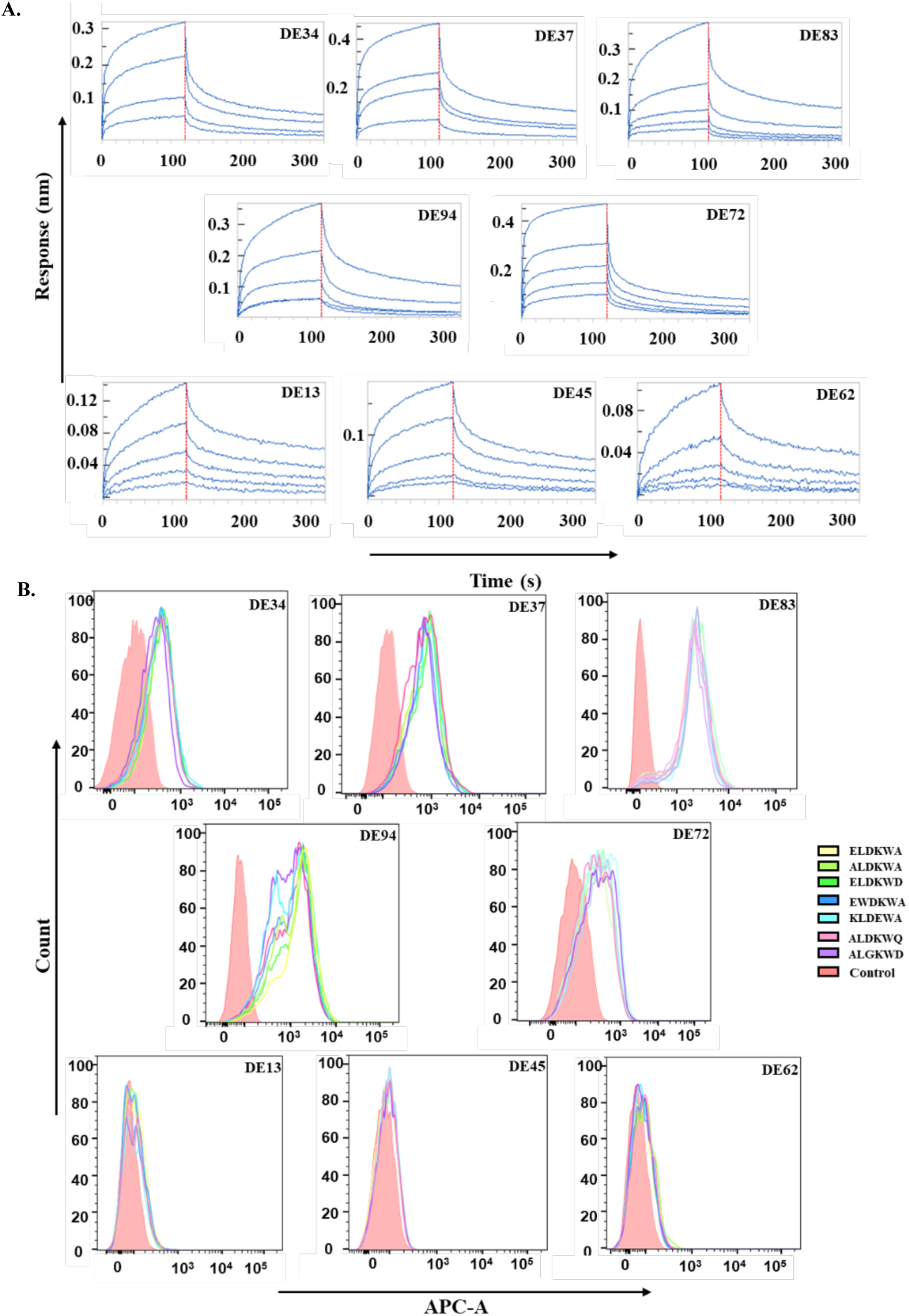
Binding analysis of the scFvs against the gp41 and gp160 proteins. **A.** BLI kinetics curve showing association and dissociation profile of scFvs against gp41 protein at different concentrations. **B.** scFvs were analyzed for their binding with the gp160 protein expressed on HEK293T cells by flow cytometry. Unstained cells were taken as a control.

To explore the multireactive potential of scFvs, a gp160 mammalian expression clone (CNE8) was expressed on HEK 293T cells, and surface expression was analyzed using flow cytometry. Variants of native epitopes were introduced gp160 mammalian plasmid using site-directed mutagenesis. Intriguingly, five out of eight scFvs showed shifts in fluorescence intensity, indicating binding to gp160 and its variants on the cell-surface (Figure 5B) and the shift was observed across all the variant tested. These results suggested that scFvs can bind the whole virus. While all scFvs bound the soluble recombinant gp41 protein, some exhibited reduced or no binding to the membrane-anchored form, likely due to limited epitope accessibility on the native-like cell surface. The binding of scFvs with all the variants unveils the multireactivity potential of these scFvs against the HIV-1 envelope protein.

### Crystallization and structure determination of scFv DE94

To get insights into the molecular interaction of scFv with peptide antigen, we attempted crystallization of all anti-GERDRDR antibodies and five anti-ELDKWA antibodies, both in apo form and with the native peptide using the hanging drop vapor diffusion method. Crystals of scFv DE94 (anti-ELDKWA antibody) were successfully obtained in the apo form that diffracted at 2.53 Å resolution and belonged to the monoclinic space group, P21, with cell dimensions of a = 43.80 Å, b = 183.92 Å, and c = 65.73 Å and β = 93.60°. Incidentally, DE94 has relatively high binding among all five anti-ELDKWA antibodies. The complete data set has Rmerge of 15.0%. Initial phasing of the scFv was performed by molecular replacement using a previously published independent scFv structure (PDB ID: 7YUE). The refined structure contains four molecules in an asymmetric unit of scFv (Figure 6A). The final Rwork and Rfree values were 19.9% and 24.3%, respectively, with 97.59% amino acid residues in the allowed region of the Ramachandran plot. All the molecules of scFv of an asymmetric unit were superimposed on each other, and no significant structural deviations were observed as evident from the RMSD values, which were in the range of 0.29-0.41 Å (Figure 6B). The electron density of chain D in DE94 was very weak. The data collection statistics and refinement statistics are shown in Table S1. The scFv DE94 comprises variable light and variable heavy chains connected via Gly-Ser linker. The six complementary determining regions (CDRs) forming the antigen-binding site are highlighted in Figure 6C. The atomic coordinates and structure factors are available in the RCSB Protein Data Bank under accession code 9V5N.

**Figure 6.**
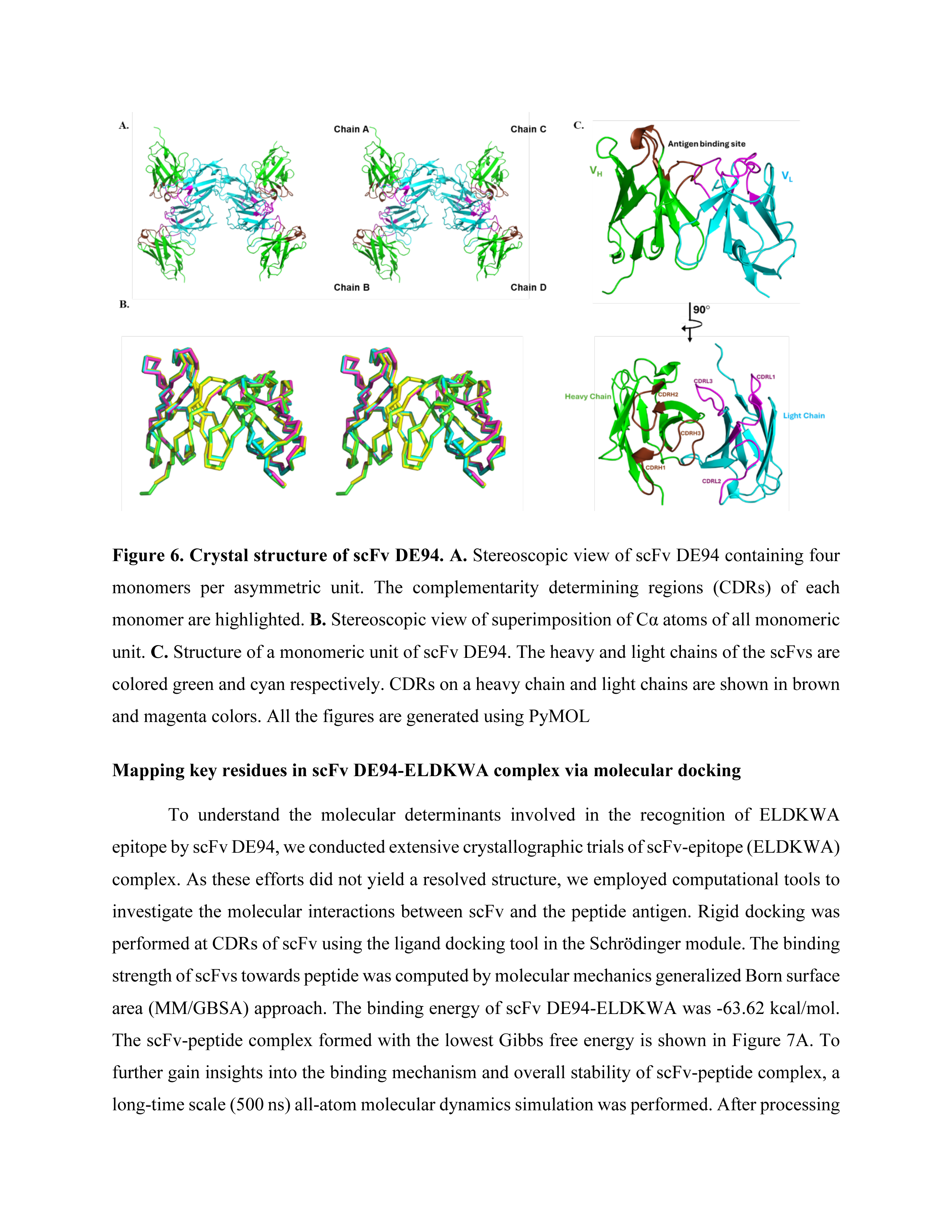
Crystal structure of scFv DE94. **A.** Stereoscopic view of scFv DE94 containing four monomers per asymmetric unit. The complementarity determining regions (CDRs) of each monomer are highlighted. **B.** Stereoscopic view of superimposition of Cα atoms of all monomeric unit. **C.** Structure of a monomeric unit of scFv DE94. The heavy and light chains of the scFvs are colored green and cyan respectively. CDRs on a heavy chain and light chains are shown in brown and magenta colors. All the figures are generated using PyMOL

### Mapping key residues in scFv DE94-ELDKWA complex via molecular docking

To understand the molecular determinants involved in the recognition of ELDKWA epitope by scFv DE94, we conducted extensive crystallographic trials of scFv-epitope (ELDKWA) complex. As these efforts did not yield a resolved structure, we employed computational tools to investigate the molecular interactions between scFv and the peptide antigen. Rigid docking was performed at CDRs of scFv using the ligand docking tool in the Schrödinger module. The binding strength of scFvs towards peptide was computed by molecular mechanics generalized Born surface area (MM/GBSA) approach. The binding energy of scFv DE94-ELDKWA was -63.62 kcal/mol. The scFv-peptide complex formed with the lowest Gibbs free energy is shown in Figure 7A. To further gain insights into the binding mechanism and overall stability of scFv-peptide complex, a long-time scale (500 ns) all-atom molecular dynamics simulation was performed. After processing the trajectories, root mean square deviations (RMSD) were examined. The system was stable throughout the run, as seen in Figure S8A. The data suggested that scFv had a stable interaction with ELDKWA, which was supported by the finding that the system’s RMSD values remained constant (Figure S9A). The backbone RMSF of the scFv was also examined throughout the 500 ns run and the Gly-Ser linker in the crystal structure connecting the heavy and light chains demonstrated high fluctuation (Figure S9B).

**Figure 7.**
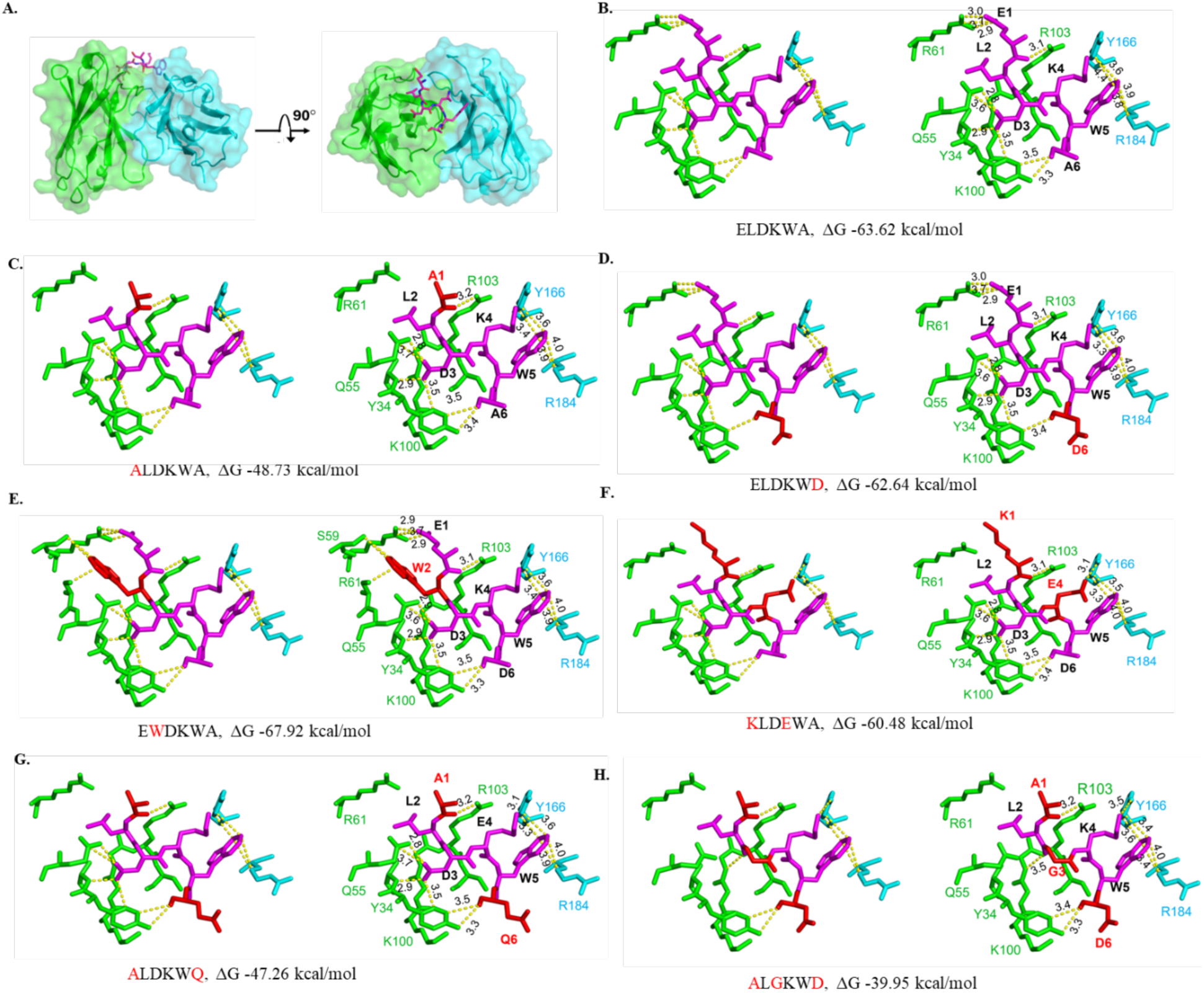
Model of scFv DE94-ELDKWA complex and stereo view of interactions with ELDKWA and its variants. **A.** Overall view of docked complex of scFv-ELDKWA where scFv is shown in surface representation and ELDKWA in stick form. **B-H**. Stereoscopic view of hydrogen bond interaction and hydrophobic interaction of scFv DE94 with ELDKWA, ALDKWA, ELDKWD, EWDKWA, KLDEWA, ALDKWQ, and ALGKWD is shown along with its Gibbs’ free energy. Ligand interaction diagrams were prepared in PyMOL with a maximum distance of hydrophobic interactions at 4.0 Å and hydrogen bonds at 3.5 Å. Heavy and light chains are represented in green and cyan, respectively, and peptides are shown in magenta.

To envisage the change in binding affinity with different variants, we mutated the native peptide within the scFv–ELDKWA complex and performed energy minimization for each mutant. The resulting models were used to calculate binding free energies using the MM-GBSA module in Schrödinger. The binding energies of all epitope variants are shown in Figure S9E. The representative structural overlays of native and mutant complexes, highlighting the modeled conformations within the binding site are shown in Figure S9 C and D. To elucidate the interaction between the scFv-peptide complex, hydrogen bond and hydrophobic interaction were analyzed. Hydrogen bonding was observed between the peptide and the residues located in the CDRH1, CDRH2, CDRH3, CDRL1, and CDRL2 regions of the scFv. In the scFv-ELDKWA complex, E1 of the peptide is involved in H-bond interaction with R61 of CDRH2 and R103 of CDRH3; D3 is involved in H-bond interaction with A35 of CDRH1 and Q55 of CDRH2, with the binding energy of -63.62 kcal/mol. While E1, D3, W5, and A7 also showed hydrophobic interaction with CDR residues as shown in Figure 7 B. All these interactions made the scFv-epitope complex stable. In the case of scFv-ALDKWA complex, binding energy has reduced to -48.73 kcal/mol, which is due to the loss of interaction of alanine with scFv. Similarly, in the case of ALDKWQ (double mutant), binding energy was -47.26 kcal/mol, as the mutation has no effect at the 6^th^ position. However, in the case of ALGKWD (triple mutant), energy has further reduced to -39.95 kcal/mol due to the loss of interaction of glycine at the 3^rd^ position. In the case of ELDKWD with the binding energy of -62.64 kcal/mol, there is no change in interaction due to the mutation at the 6^th^ position. In double mutant KLDEWA, the binding energy is equivalent to the native epitope as the mutation from E to K at the first position is balanced by the mutation from K to E at 4^th^ position. EWDKWA has more negative ΔG (-67.92 kcal/mol) showing a more stable complex which can be verified by the additional interaction of W4 with CDRs of scFv (Figure 7C-H). Although there was a difference in the binding energy of the native epitope with its variants, it was noteworthy that ΔG for all the epitopes was in the range of antigen-antibody interactions.

## Discussion

The monoclonal antibodies are recognized as a key mediator of protective immunity against HIV-1 [30, 31]. There is a need to develop broadly cross-reactive therapeutic antibodies. While cross-reactive antibodies are known to exist in the immune system, immune evasion in HIV is also well-documented. This study aimed to explore antibody cross-reactivity and immune escape mechanisms by comparing antibodies against two critical epitopes of the HIV envelope protein gp41: GERDRDR and ELDKWA and to draw insights from these findings in the broader context of other fast-mutating viruses.

Antibodies targeting the epitope GERDRDR exhibited greater cross-reactivity compared to those against ELDKWA, reflecting different hotspot behaviors. Together, both epitopes demonstrated structural plasticity in antigen-combining sites. The broader reactivity observed against GERDRDR-specific epitopes can likely be attributed to multiple charged amino acids across the epitope sequence, enabling the binding of scFvs to all the variants. In contrast, antibodies against ELDKWA showed less cross-reactivity, only five out of eight binders could recognize the native epitope and its variants. A plausible explanation for the lack of binding by the remaining three scFvs could be steric hindrance from the membrane-anchored protein.

Notably, during HIV-1 infection, polyreactivity is observed in approximately 70–75% of anti-gp160 antibodies, and this property is even more pronounced among anti-gp41 antibodies, of which 85–90% exhibit polyreactivity [22, 29, 32]. This high proportion of polyreactivity highlights the significance of multispecific antibodies in shaping the immune response. Moreover, our findings support the hypothesis that B cell clones producing multispecific antibodies undergo positive selection during the affinity maturation [33]. This is in line with the previous studies indicating that the primary factor contributing to antibody multispecificity in response to HIV Env proteins is the inherent conformational flexibility of the antigen-binding site [34, 35]. This structural plasticity is not unique to HIV but also observed in other rapidly mutating viruses. For instance, multispecific antibodies have been identified in the influenza virus, against a key antigenic target hemagglutinin (HA) protein [5]. The HA protein undergoes significant antigenic drift leading to the emergence of new strains. The presence of multispecific antibodies in the humoral immune response would be necessary for preventing immune evasion during influenza infections despite mutations that are beneficial for viral escape and lead to antigenic diversification [5]. This phenomenon is similar to what we observed in the case of HIV, where antibodies with broad reactivity can accommodate diverse variants due to the structural flexibility of the epitopes. Similarly, in SARS-CoV-2, multi-specific antibodies have been identified against the spike protein. Antibodies produced following vaccination against SARS-CoV-2 have displayed the capability to neutralize a range of emerging strains, which may be attributed to the multi-specific nature [4, 36]. The broad reactivity of such antibodies is crucial for effective immunity, especially as new variants continue to emerge.

Our findings correlate with previous findings that the immune system can produce antibodies with degenerate potential, enabling recognition of the cross-clade variants [34, 35]. The structural plasticity observed in both gp41 epitopes highlights the adaptability of multispecific antibodies, which is crucial for combating HIV’s high mutation rate and ensuring an effective immune response. To gain deeper insights into molecular interactions, we performed crystallographic studies on scFv DE94. These studies, combined with in-silico analysis of scFv DE94 binding to ELDKWA and its mutants, revealed that the binding energy ranged from −39.95 to −67.92 kcal mol^−1^, indicating strong interactions, however certain mutations led to reduced binding energy. These results support our hypothesis that multispecific antibodies can accommodate antigenic diversity, although certain mutations may impact binding efficacy.

Previous studies have reported that neutralizing antibodies targeting the MPER of HIV-1 gp41 possess cross-reactivity [7, 8]. Specifically, the broadly neutralizing antibody 2F5, which targets the ELDKWA epitope relies on DKW residues for interaction [27] and substitution from DKW to DSW makes the virus often resistant to antibody neutralization [37]. In the present study, the scFvs demonstrated remarkable recognition of mutations in the DKW region, except for scFv DE62, which failed to recognize the mutant peptide KLDEWA, and scFv DE72, which exhibited an inability to recognize the triple mutant peptide ALGKWD as confirmed through ELISA and BLI assays. Nevertheless, when the variants were exposed on the surface of mammalian cells, cross-reactivity was observed with all the variants. This observation may be attributed to conformational constraints imposed by the membrane-anchored protein.

Our observations are similar to findings in other rapidly mutating viruses, such as influenza and SARS-CoV-2, where multispecific antibodies are crucial in neutralizing diverse strains despite antigenic drift [4, 36]. HIV exhibits a high mutation rate with approximately 10^−3^ substitutions per nucleotide per replication cycle, contributing to its genetic diversity [38, 39]. However, despite this, its evolution is comparatively slow at the population level [6–8]. This could be due to number of factors such as less selection pressure over time, weak immune response, non-beneficial mutations affecting viral fitness, reversal of patient-specific adaptive changes following transmission and retrieval mechanism in which initially infected virus is preferentially transmitted [41, 42]. The presence of multispecific antibodies may help in constraining the evolution of new strains by providing broad protection against diverse strains. Our finding suggests that not all mutations lead to antigenic drift, only crucial mutations significantly alter viral antigens, rendering existing antibodies less effective.

In conclusion, this study highlights the importance of antibody cross-reactivity and structural plasticity in HIV research. The high prevalence of cross-reactive antibodies and their multispecific nature emphasize the need for vaccines and therapeutics that can accommodate viral diversity. Insights gained from crystallographic studies and antibody behavior, particularly regarding the impact of mutations, are valuable for developing novel strategies to enhance protection against mutant strains of viruses and improve disease control efforts.

## Methods

### Peptide selection, synthesis, and conjugation to BSA

The neutralizing epitopes were selected from the HIV gp41 region. After selecting two critical epitopes from MPER and CTT regions, escape variants of both epitopes were chosen from the HIV sequence database (http://www.hiv.lanl.gov/). We have analyzed around 12000 variants of these epitopes present in the database, resulting in the selection of six variants against each epitope. All the peptides were synthesized with N-terminal cysteine/lysine for BSA conjugation using an automated peptide synthesizer (433A; Applied Biosystems, Foster City, CA, USA). The crude peptides were purified by reverse-phase semi-preparative HPLC (Waters, USA) using a C18 column. Conjugation of the peptide with bovine serum albumin (BSA) was done either using lysine or cysteine residue of the peptide. The conjugation using lysine residues was done following the glutaraldehyde cross-linking method whereas cysteine-based conjugation was done using N - succinimidyl 3-(2-pyridyldithio) propionate (SPDP).

### Biopanning

Tomlinson I and Tomlinson J, antibody phage display libraries, KM13 helper phage and *Escherichia coli (E. coli)* TG1-Tr and HB2151 were procured from Centre for Protein Engineering, MRC Laboratories, Cambridge, United Kingdom while HuScL4 libraries were procured from Creative Biolabs, New York, USA. For screening of libraries, MaxiSorp or PolySorp immunotubes (in every alternate round) were immobilized with 4 ml of 100 μg/mL BSA-peptide and BSA in 1x PBS and kept at 4°C overnight. Thereafter, the tubes were washed three times with 1x PBS and blocking was done with 2% skimmed milk in 1x PBS for 2 hours at room temperature (RT). To remove BSA-specific phages, around 10^12^ phages of amplified Tomlinson I, J, and HuScL4 libraries or amplified phages (from successive rounds of selection) were pre-incubated with BSA-coated immunotubes for 1 hour, and then the remaining phages were incubated with BSA-peptide for 1 hr on a rotating platform and 1 hour of static incubation. The unbound phages were removed, and washing was done with 1x PBS with 0.05% Tween 20 (10 times for selection round 1 and incremented 10 times after every subsequent round) to remove the weakly bound phages. Affinity elution was done with 0.45 mL of 1 mg/ml of peptide for 30 min to elute the bound phages [5]. The eluted phages were then treated with 50 μl of 10 mg/mL bovine trypsin (Type XIII from Bovine Pancreas, Sigma-Aldrich, St. Louis, MO) for 10 min. The eluted phages were further amplified and used as input phages for the next round. The biopanning was repeated 4-5 times so that the high-affinity phages bind to the peptide epitope.

### Polyclonal and monoclonal phage ELISA

The phages obtained after 4 or 5 rounds of selection were checked for their binding towards the peptide epitope. A 96-well plate was coated with 100 μg/mL of peptide-BSA conjugate and BSA. The following day, the plates were washed thrice with 1x PBS and blocked with 2% skimmed milk in 1x PBS for 2 hr. The polyclonal phages/monoclonal phages were added to the antigen-coated plates and incubated for one hour at RT [40]. The phages were removed, and plates were washed thoroughly with 0.1% Tween 20 in PBS (PBST). The anti-mouse M13-HRP was added at 1:5000 dilution for 1hr at room temperature. The plate was washed 4 times with PBST and subsequently developed with ortho-phenylenediamine (OPD) and H2O2 as substrate and the absorbance was read at 490 nm.

### Cloning, Expression and purification of scFvs

The selected scFvs from Tomlinson libraries were cloned into pET22b vector, while scFvs from HuScL4 library were cloned into pET22 b-DS55 (Addgene ID-187175). In the commercially available pET22b vector (Addgene-Catalog number 69744-3), the Sal-I and Hind-III sites were followed by Not-I, Xho-I, and 6X-His tags. We have modified this vector such that Hind-III site is followed by Sal-I, Not-I, Xho-I and 6X-His tag to express scFvs from HuScL4 library in-frame. The amber stop codon present in the scFvs was mutated to glutamine by site-directed mutagenesis. For overexpression of scFvs, clones were transformed into the BL21 *E. coli* strain. The culture was inoculated and incubated at 37°C for 5 hr, followed by induction with Isopropyl ß-D-1-thiogalactopyranoside (IPTG) and incubated at 18°C for 16-18 hr. The pellet obtained was resuspended in periplasmic extraction buffer I (100mM Tris pH=8, 20% sucrose. 1mM EDTA) and incubated on ice for 45 min. After centrifugation, the pellet was dissolved in periplasmic buffer II (5 mM MgCl2) and incubated on ice for 30 min. After centrifugation, supernatant from both processes was mixed and used for immobilized metal affinity chromatography (IMAC). Further size exclusion chromatography (SEC) was performed using a HiLoad 16/60 Superdex 75 prep grade column by fast performance liquid chromatography (ÄKTA pure) at 4°C.

### Titration ELISA

A 96-well ELISA plate was coated with 10 µg/ml of BSA-peptide and BSA in 1x PBS overnight at 4°C. BSA was coated as a control. After 1x PBS washing, plates were blocked with 2% skim milk in 1x PBS for two hours at room temperature. Purified antibodies were titrated from 50 µg/ml to 16 dilutions and incubated for 1 hour at 37°C. After three washes with PBST, the plates were incubated for 1 hour at 37 °C with a 1:5000 dilution of an anti-His HRP-conjugated antibody (Santa Cruz Biotechnology, Cat# sc-8036). Plates were again washed three times with PBST solution. Color development was achieved using substrate o-phenylenediamine (OPD; HiMedia) in the presence of hydrogen peroxide (H₂O₂; Sigma-Aldrich). The optical density (OD) was measured at 490 nm using a SpectraMax plate reader (Molecular Devices).. EC50 were calculated with GraphPad Prism version 6.01.

### Bio-layer interferometry

Binding of scFvs with peptide as well as protein was done by using Bio-layer interferometry (BLI), Octet RED96e instrument (ForteBio, Molecular Dimensions). Streptavidin (SA) biosensors were used to immobilize biotinylated BSA-peptide/gp41 and BSA as a control till it reached a nanometer shift of 0.8-1.2nm. The scFv was prepared in 1x PBS in two-fold dilution for affinity analysis. The regeneration and neutralization buffers used during the experiment are 10 mM glycine, pH = 2.5, and 1x PBS, respectively. The association and dissociation steps were recorded for the 120s and 200s, respectively. The data were analyzed using Forte Bio Data analysis software 10.0.0.1. The global data specifying the fitting to a 1:1 binding model was used to determine the kon, koff and KD.

### Flow cytometric binding assay and site-directed mutagenesis

The scFv phage clones displaying premature termination codon amber (G68, G73, G77, and G80) were mutated to glutamine by site-directed mutagenesis for expression in the BL21 strain. To analyze the binding of scFvs to the full-length gp160 protein, flow cytometry experiments were done using the HEK293T cells. HIV full-length gp160 expressing clones, named CNE8 and SF162, were procured through the NIH AIDS reagent program. Epitope analogs were generated via site-directed mutagenesis using QuickXL-II SDM kit (Agilent Technologies-Catalog #200521) as per the manufacturer’s protocol.

Human embryonic kidney 293T cells (HEK293T; procured from ATCC) were seeded in a 60 mm dish in DMEM containing 10% fetal bovine serum (FBS), 1% penicillin, and streptomycin antibiotics. The next day, cells with a density of 50-60% were transfected using PEI 25K reagent (Polysciences-Catalog #23966) in OptiMEM media. PEI was mixed with plasmid DNA in 3:1 ratio and kept at room temperature for 30 min. DNA and PEI complex were added slowly into the culture dish and kept for 4-6 hr at 37°C in a 5% CO2 incubator. The media was changed with the complete media and incubated for 36 hours. The cells were washed to make a single-cell suspension and blocked with 2% FBS in 1x PBS (FACS buffer) for 1 hour at 4°C. Cells were fixed using BioLegend Fixation buffer (Catalog-#420801) and permeabilized using Intracellular Staining Perm Wash Buffer (BioLegend-Catalog #421002) to access the GERDRDR epitope inside the viral membrane. After that, cells were incubated with scFvs for 1 hr at 4°C and washed thrice with FACS buffer. His-Tag (D3I1O) XP^®^ Rabbit mAb (Alexa Fluor^®^ 647 Conjugate, CST Cat# 14931) was added as a secondary antibody at the dilution of 1:250 for 20 min at RT. Cells were washed thrice with FACS buffer and resuspended in 430 μl 1x PBS, and samples were acquired in BD LSRFortessa using Diva software (BD Bioscience).

### X-ray crystallography and structure determination

Crystallization of scFv DE94 was performed using hanging drop vapor diffusion method using Mosquito LCP nano-dispenser (TTP Labtech). Crystals were grown in conditions containing 0.5 M SPG buffer, pH=4, 25% PEG1500, 10% glycerol. The data were collected at ID30B beamline, European Synchrotron Radiation Facility (ESRF) with 15 % glycerol as cryoprotectant. Following data collection, HKL3000 software was used to process the data. PHASER from the CCP4 program suite was used to determine the structure by molecular replacement with the previously published structure of scFvs (PDB: 7YUE) as a starting model. Refinement was carried out in PHENIX, and Coot was used for visualization. All the figures were prepared using PyMOL and Coot. The PDB validation server checked the final quality of the structure.

### Molecular docking and MD simulations

The scFv-peptide complex was generated by using ligand-docking tool in Schrödinger Suite [41]. The crystal structure of scFv obtained was taken as a reference. The complex was subjected to a molecular dynamics simulation run using the Desmond 3.1 MD package (Schrödinger Inc.) [42]. Mutants of epitope were generated in PyMOL, and energy minimization of all the complexes was done to check any steric hindrance in the complexes, and then Prime MM-GBSA was subjected to run. The relative binding affinity of the epitopes was calculated using the Prime MMGBSA module of Schrödinger. The ligand interaction diagram was prepared in PyMOL for interaction studies.

## ACCESSION NUMBERS

The atomic coordinates and structure factors have been deposited in the Protein Data Bank, www.rcsb.org (**PDB: 9V5N**).

## ACKNOWLEDGEMENTS

We thank Mr. Romain Talon for providing support with data collection at the European Synchrotron Radiation Facility (ESRF), Grenoble, France. We would like to thank Mr. Ravinder Kumar for assisting in cell culture. We would like to acknowledge the Department of Biotechnology, Govt. of India for the generous funding.

## AUTHOR CONTRIBUTION

D.M.S. provided supervision of the project; D.J., and D.M.S. conceived and designed the experiments; D.J. G.K. and K.J.K. synthesized the peptides. D.J. and S.V. carried out library screening. D.J. carried out scFv purification, ELISA, BLI, FACS, crystallization, structure determination and MD simulation; D.J. and Z.K.M carried out structure refinement. D.J. wrote the original draft; D.M.S. and D.J. reviewed and edited the draft and analyzed the data. All authors reviewed and approved the final version of the manuscript.

## Supplementary figures

**Figure S1.**
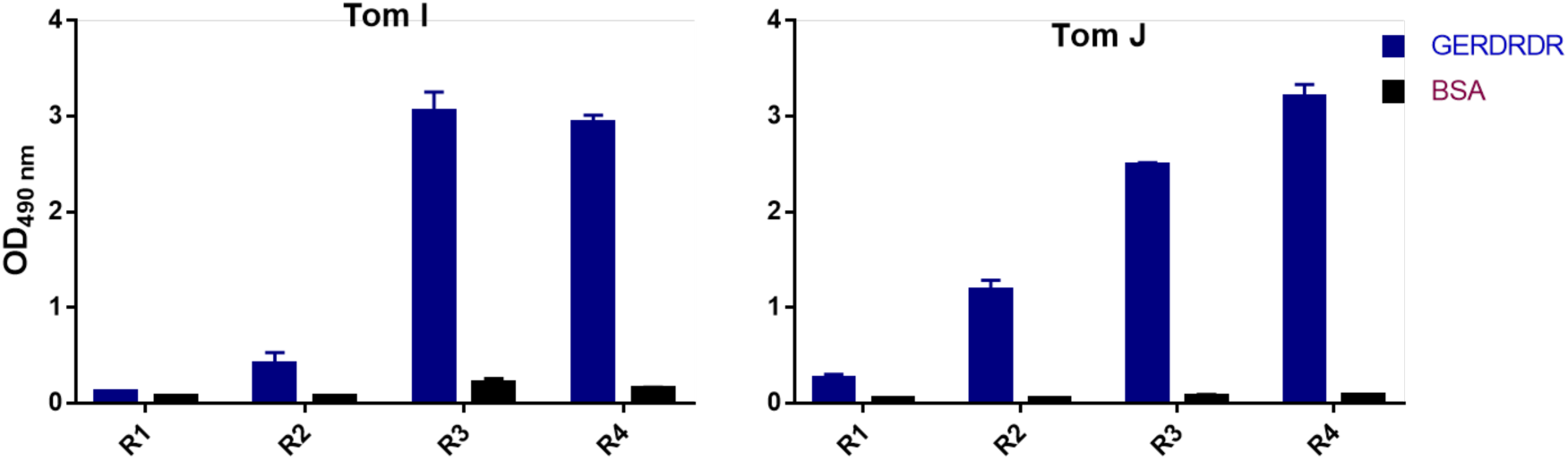
Binding of polyclonal phages against GERDRDR by ELISA. Amplified phages from each round of biopanning were used to check binding against GERDRDR using human-based scFv library.

**Figure S2.**
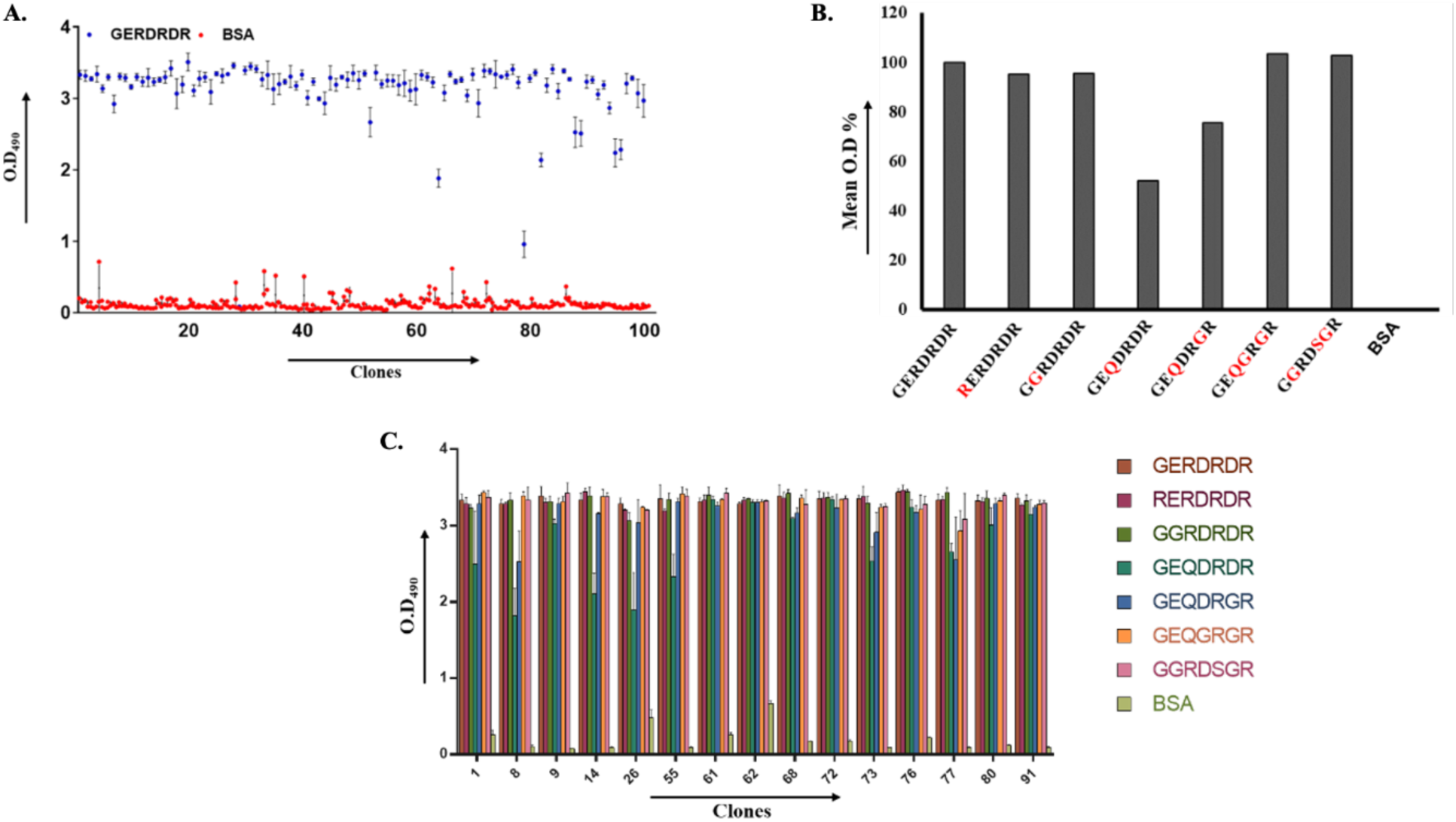
Selection and Binding Analysis of Cross-Reactive Phage Clones Targeting GERDRDR. **A.** Selection of high binding monoclonal phages from the 384 isolated phages against GERDRDR. Error bar represents standard deviation from the 3 replicates **B.** Analysis of mean O.D. percentage of scFv phages against native epitope and all the variants showing cross-reactivity. The mean O.D value percentage is 100% for native epitope and 0% for BSA. **C.** Binding of selected 15 phage clones against all the variants of GERDRDR by ELISA. Data was plotted from the previous 98 phage clones showing cross reactivity from Figure. 2B

**Figure S3.**
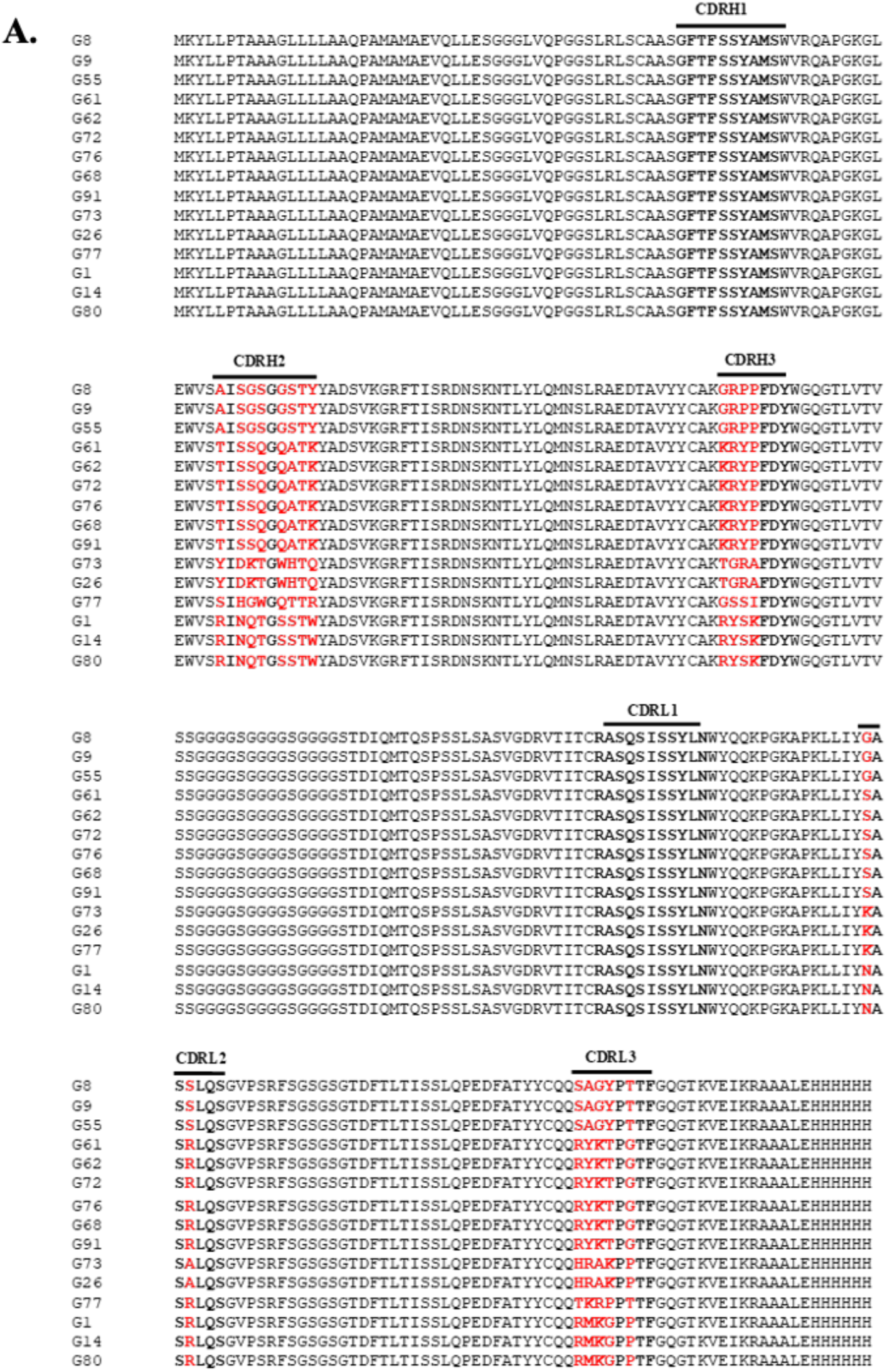
Sequence analysis of selected anti-GERDRDR clones. Multiple sequence alignment of selected 15 high binders showing diversity in CDR regions.

**Figure S4.**
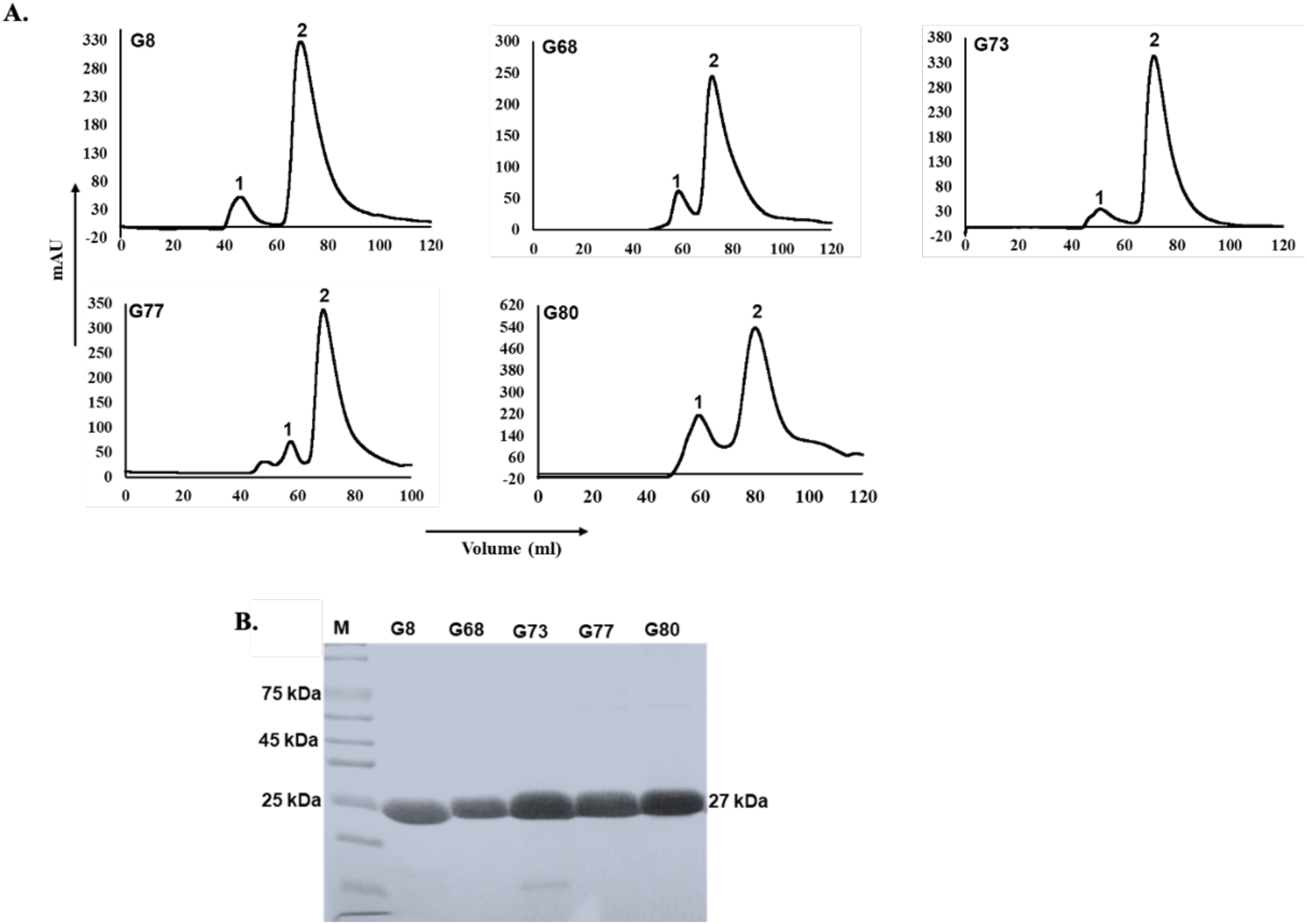
Purification profile of scFvs. **A.** Chromatogram representing purification of each scFvs by gel filtration chromatography. Peak 1 corresponds to void volume and peak 2 represents scFv eluted at 70 ml retention volume. **B.** Coomassie stained SDS-PAGE gel displaying peak 2 for each case representing scFv at 27 kDa.

**Figure S5.**
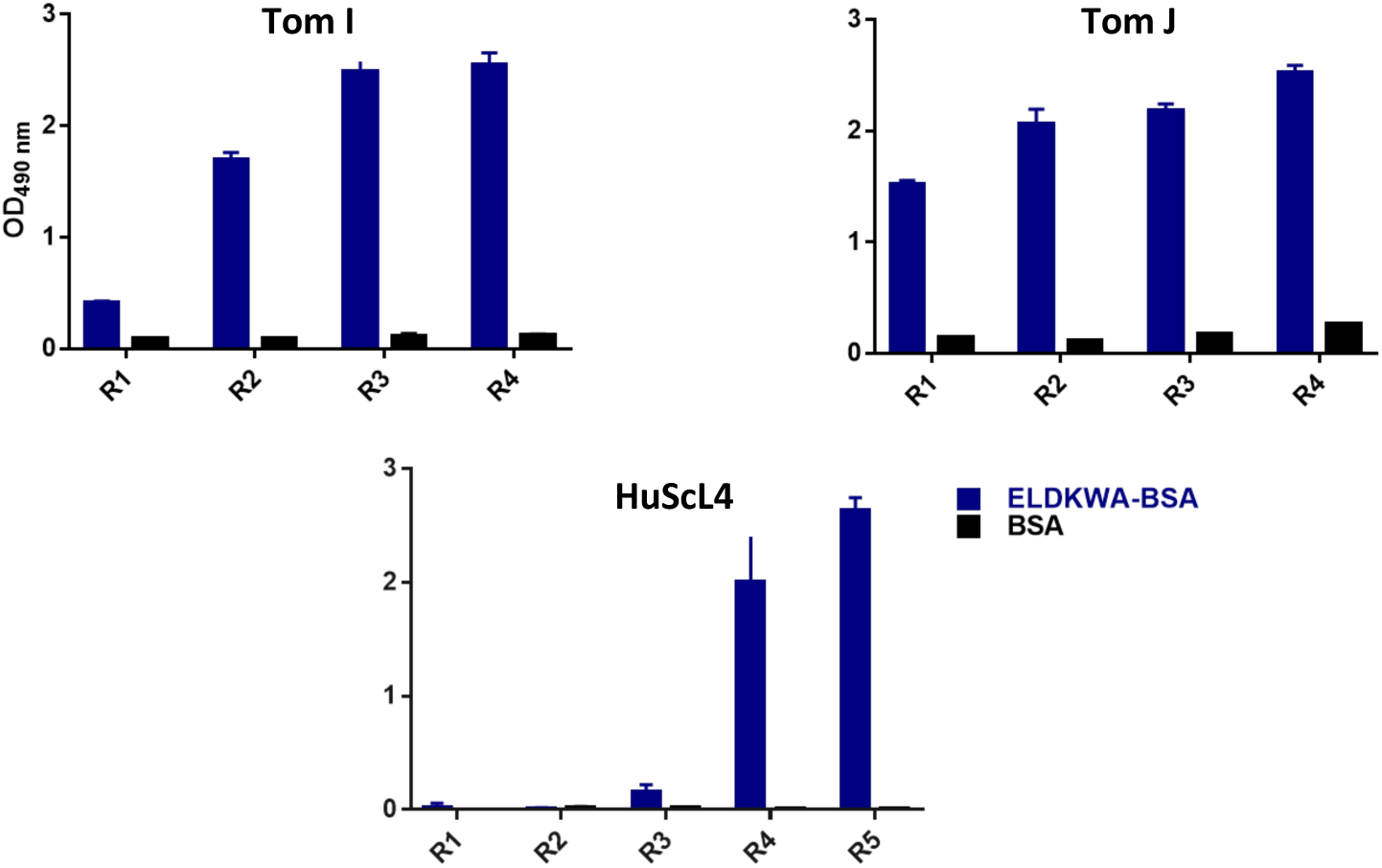
Binding of polyclonal phages against ELDKWA by ELISA. Amplified phages from each round of biopanning was used to check binding against ELDKWA using human-based scFv library.

**Figure S6.**
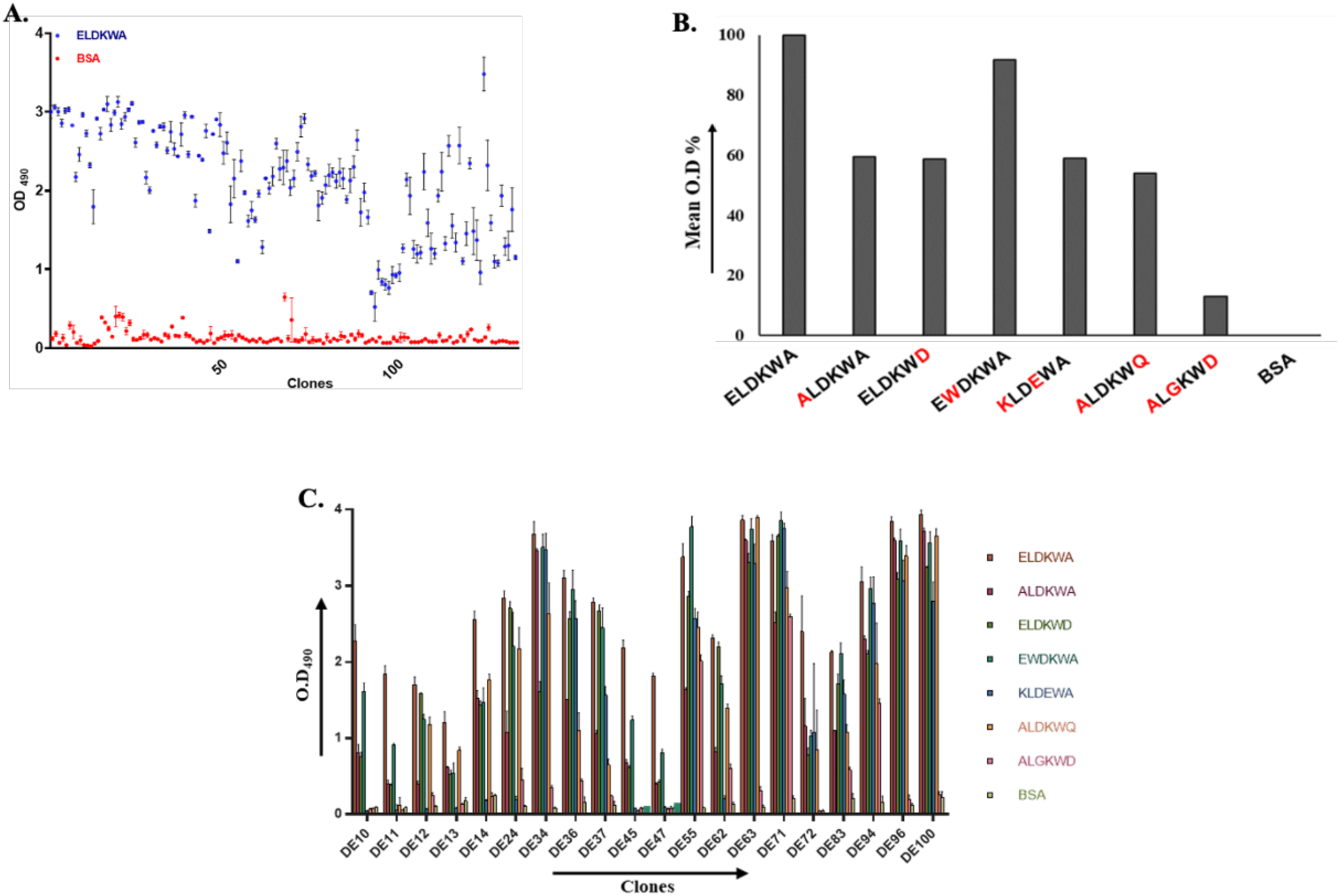
Selection and Binding Analysis of Cross-Reactive Phage Clones Targeting ELDKWA. **A.** Selection of high binding monoclonal phages from the isolated 1150 phages against ELDKWA. Error bar represents standard deviation from the 3 replicates **B.** Analysis of mean O.D. percentage of scFv phages against native epitope and all the variants showing cross-reactivity**. C.** Binding of selected 20 phage clones against all the variants of ELDKWA by ELISA. Data was plotted from the previous 80 phage clones showing cross reactivity from Figure. 4B.

**Figure S7.**
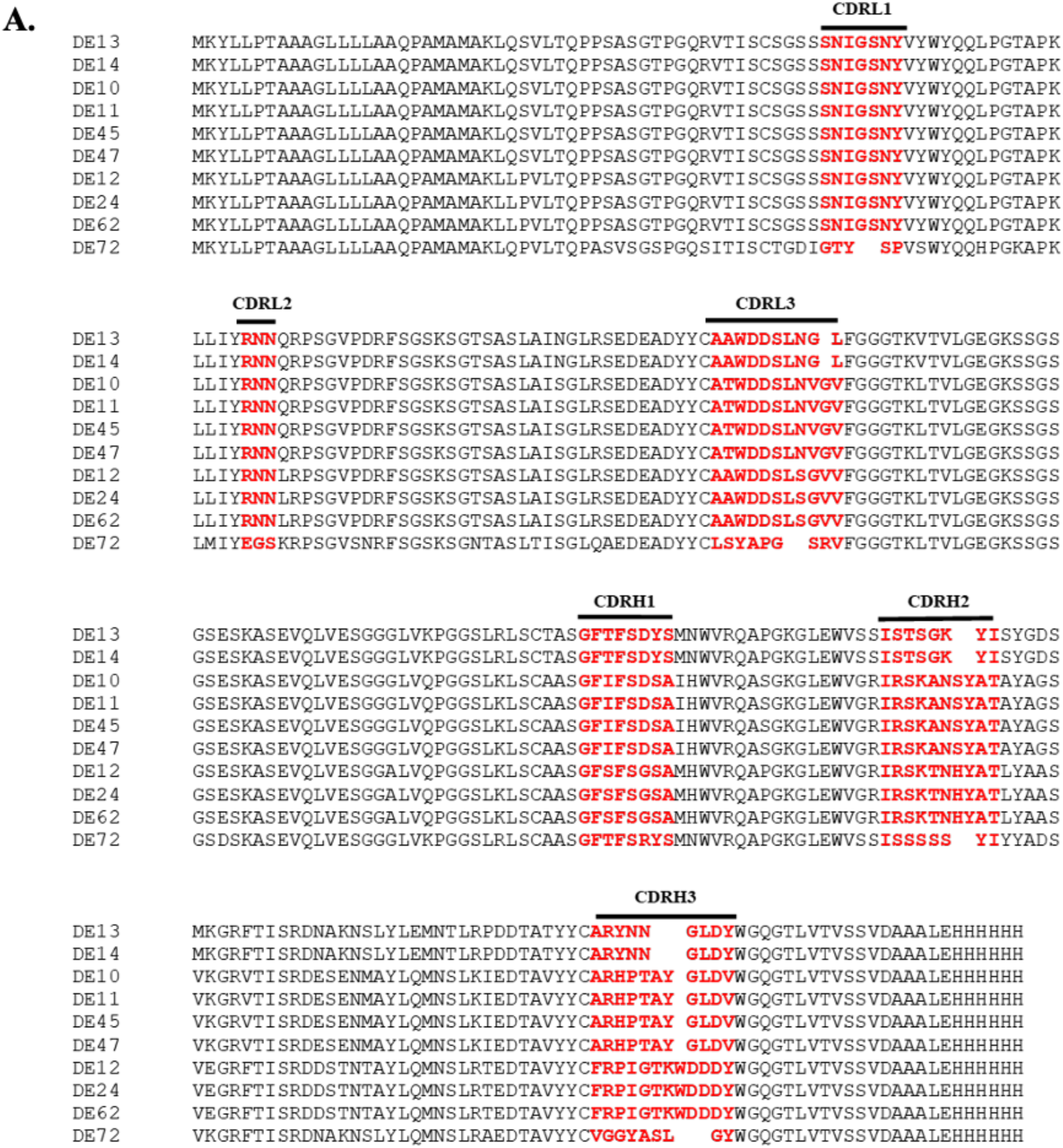
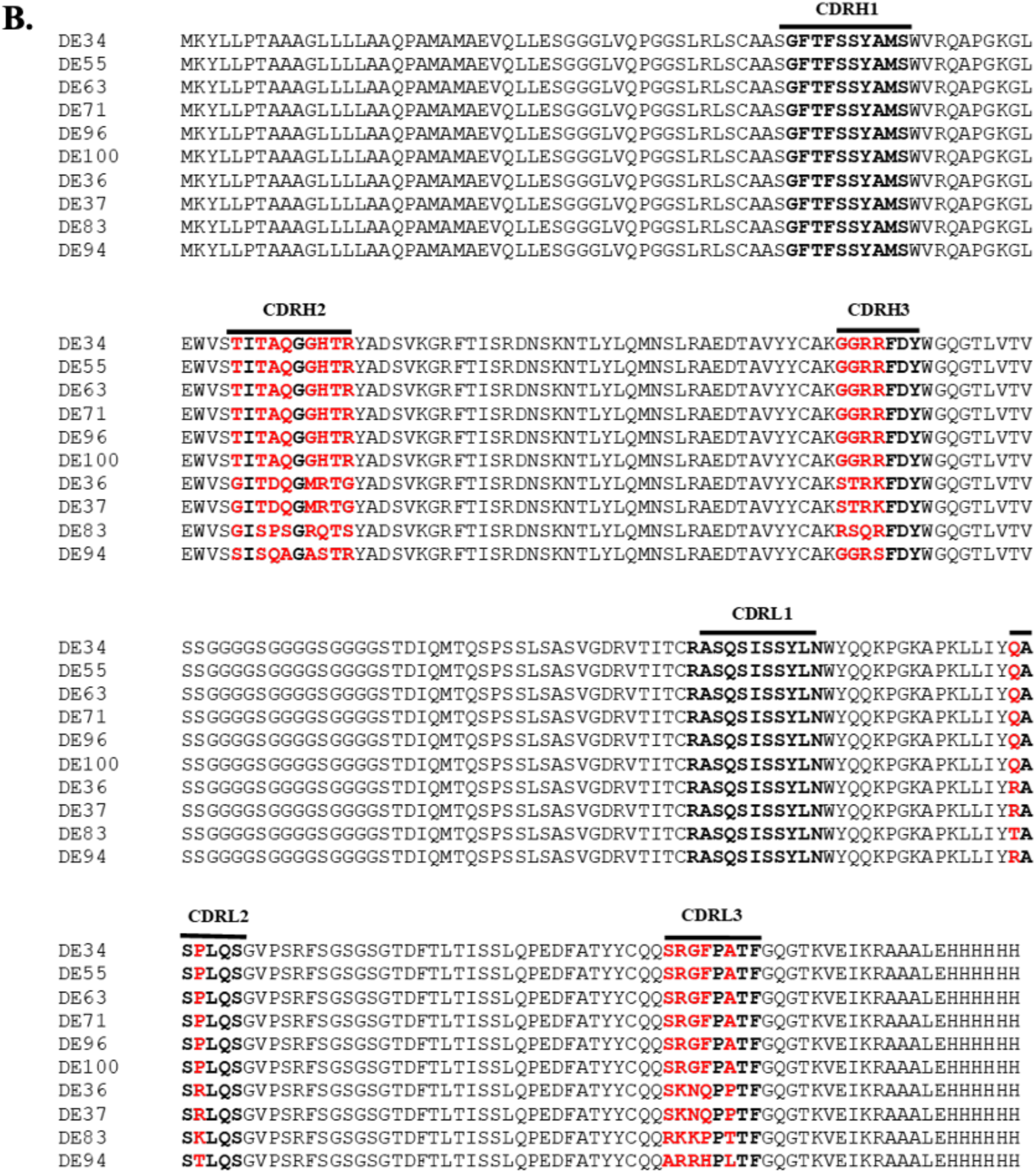
Multiple sequence alignment of anti-ELDKWA scFvs. Sequence of all the scFvs were aligned using Clustal omega. Amino acid difference in CDRs of scFv are highlighted in red and consensus sequence are shown in black. **A.** Ten scFvs are in VL-VH format while **B.** other ten scFvs are in VH-VL format.

**Figure S8.**
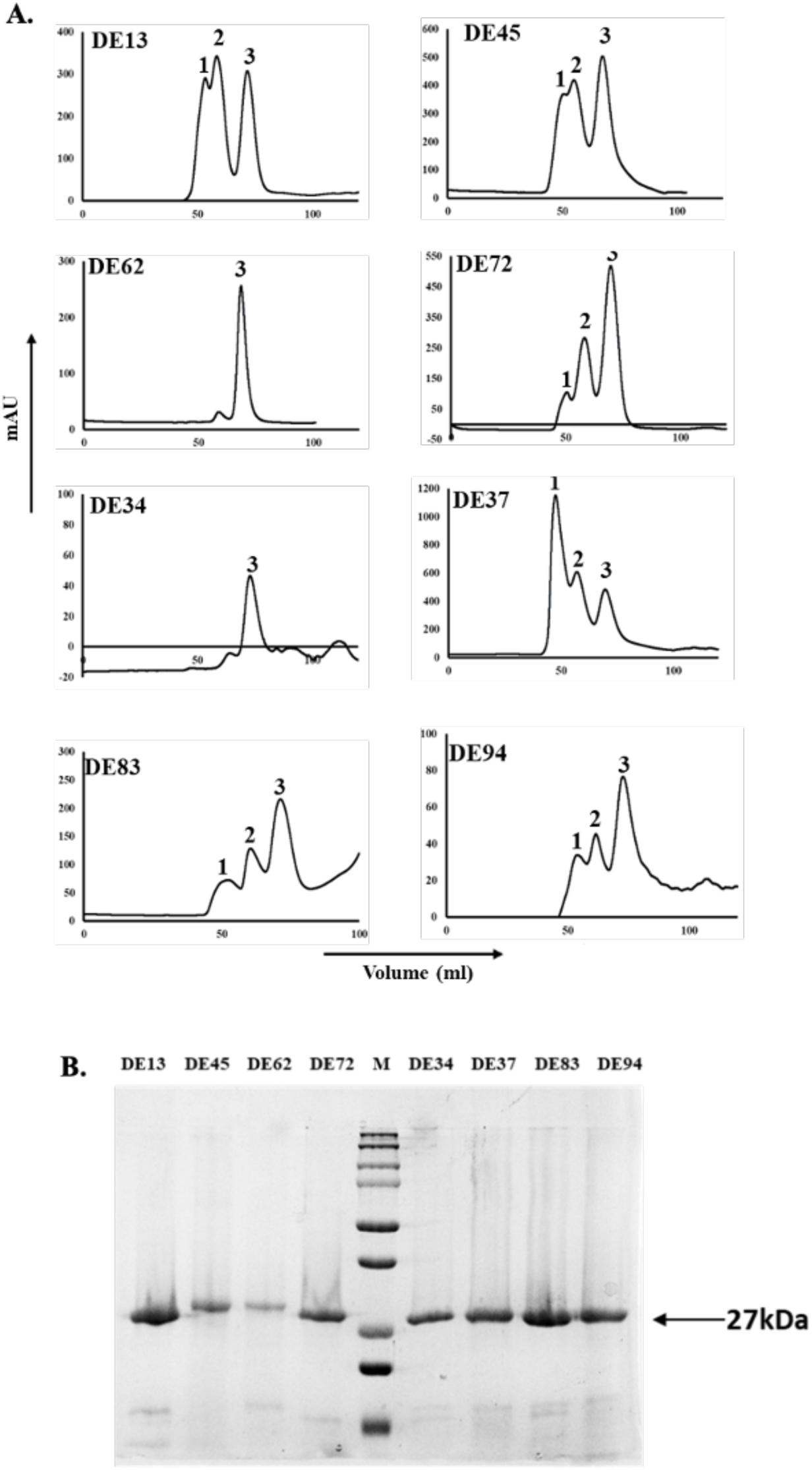
Purification profile of scFvs. **A.** Gel filtration chromatography profile of the scFvs. Peak 1 represents the column void volume, peak 2 corresponds to oligomeric form of scFv and peak 3 represents scFv monomeric form at retention volume of 70-75 ml. **B.** Coomassie stained SDS-PAGE gel displaying peak 3 for each case representing scFv at ̴27 kDa.

**Figure S9.**
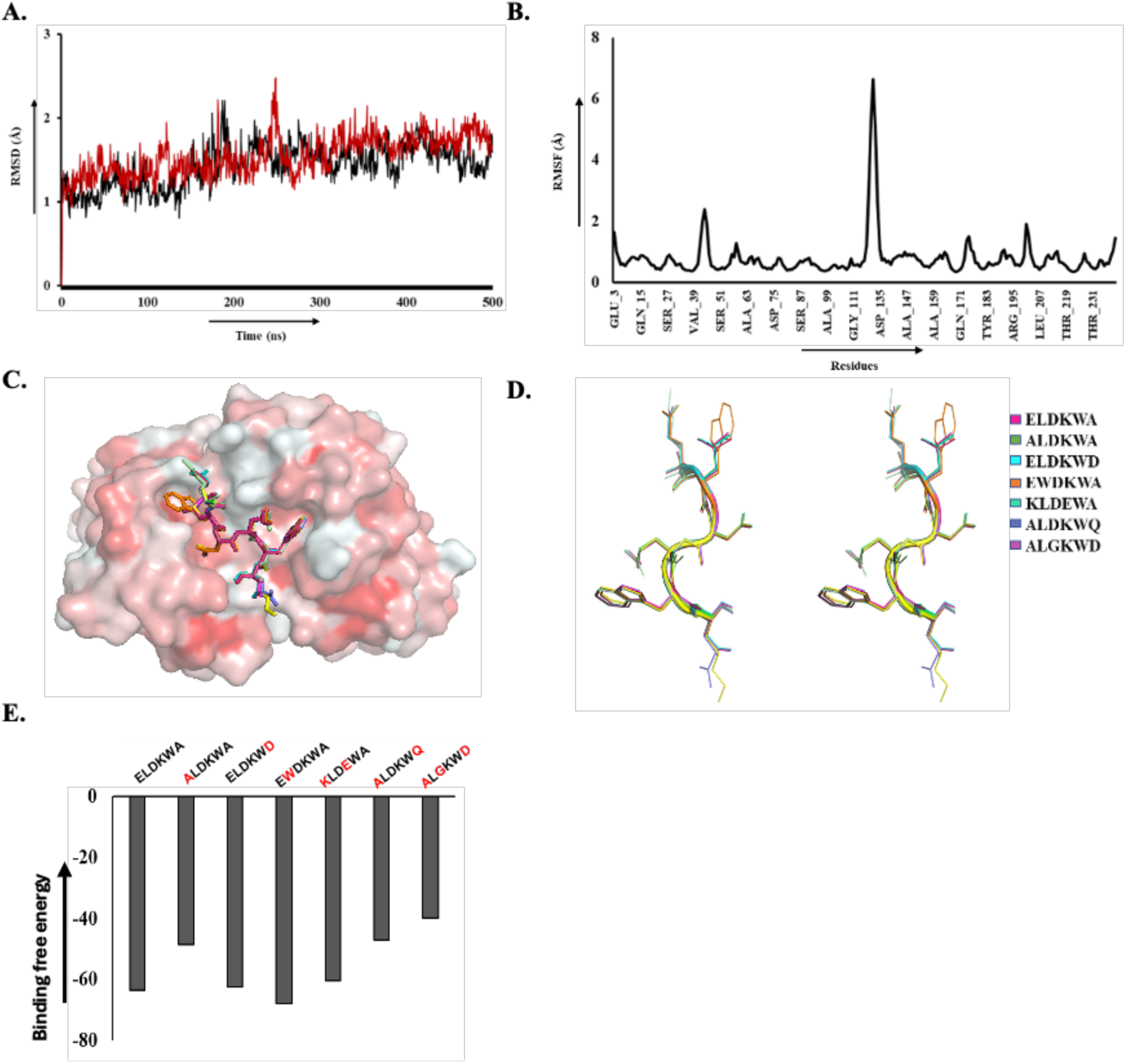
MD simulation and superimposition of scFv with native epitope and all the mutants. **A.** RMSD vs time plot of scFv-epitope complex shown in black and scFv alone in red. **B.** RMSF of scFv backbone in complex. **C.** All the complexes were superimposed showing surface representation of scFv and epitopes are shown in stick form. **D.** Stereoscopic representation of superimposed native epitope and all the variants in antibody bound confirmation. **E.** Relative binding affinity of native and seven mutant complexes calculated by Prime MMGBSA approach of Schrödinger suite. Native epitope mutants were generated using PyMOL and each complex was energy minimized.

**Table S1.**
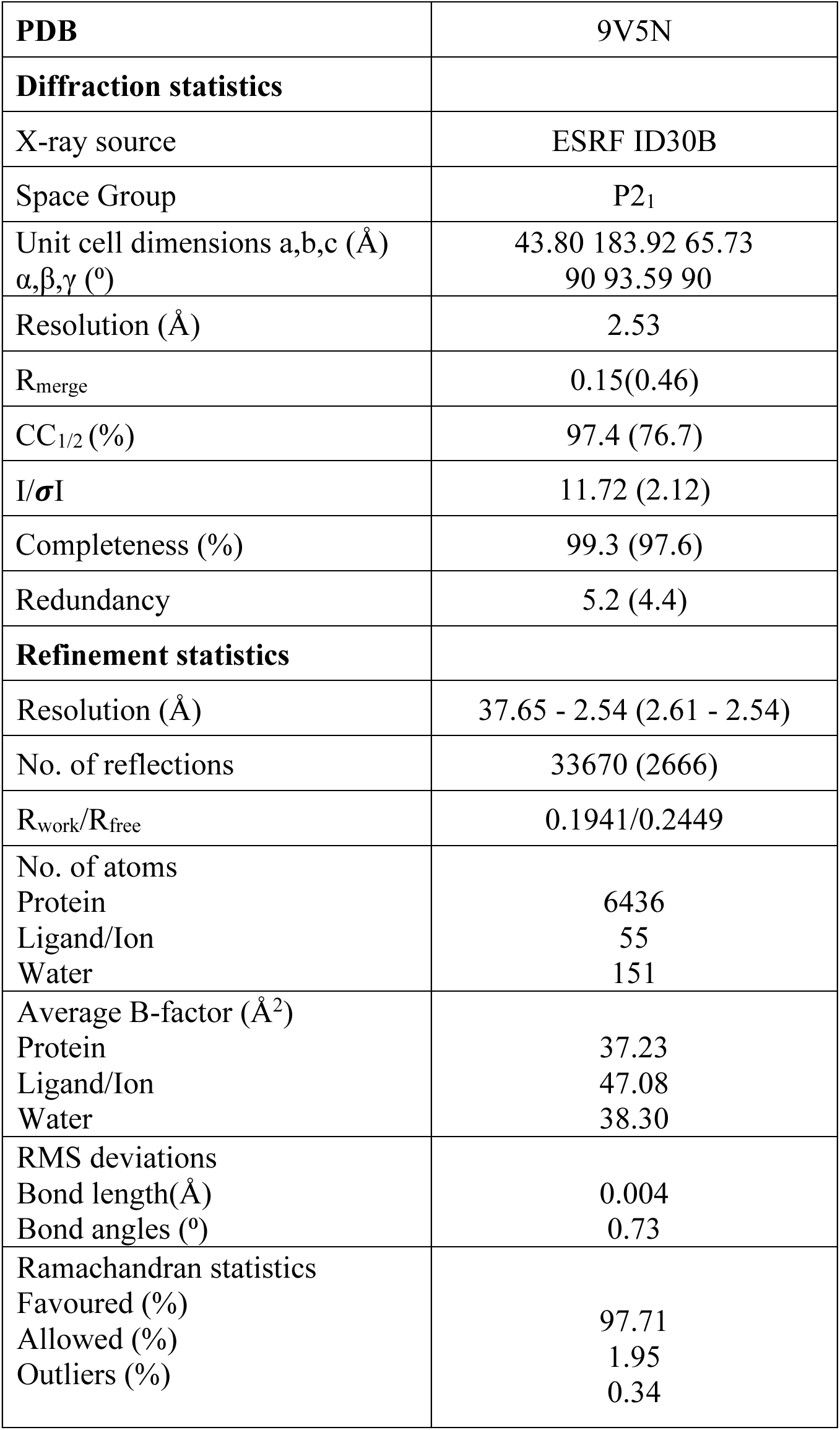
Data collection and data refinement statistics. Values in parentheses are for the highest resolution shell.

|  |  |
| --- | --- |
| **PDB** | 9V5N |
| **Diffraction statistics** |  |
| X-ray source | ESRF ID30B |
| Space Group | P2<sub>1</sub> |
| Unit cell dimensions a,b,c (Å)<br>$\alpha,\beta,\gamma$ (°) | 43.80 183.92 65.73<br>90 93.59 90 |
| Resolution (Å) | 2.53 |
| R<sub>merge</sub> | 0.15(0.46) |
| CC<sub>1/2</sub> (%) | 97.4 (76.7) |
| I/σI | 11.72 (2.12) |
| Completeness (%) | 99.3 (97.6) |
| Redundancy | 5.2 (4.4) |
| **Refinement statistics** |  |
| Resolution (Å) | 37.65 - 2.54 (2.61 - 2.54) |
| No. of reflections | 33670 (2666) |
| R<sub>work</sub>/R<sub>free</sub> | 0.1941/0.2449 |
| No. of atoms |  |
| Protein | 6436 |
| Ligand/Ion | 55 |
| Water | 151 |
| Average B-factor (Å<sup>2</sup>) |  |
| Protein | 37.23 |
| Ligand/Ion | 47.08 |
| Water | 38.30 |
| RMS deviations |  |
| Bond length(Å) | 0.004 |
| Bond angles (°) | 0.73 |
| Ramachandran statistics |  |
| Favoured (%) | 97.71 |
| Allowed (%) | 1.95 |
| Outliers (%) | 0.34 |

## References

1. Amit AG, Mariuzza RA, Phillips SE, Poljak RJ: Three-dimensional structure of an antigen-antibody complex at 2.8 A resolution. Science 1986, 233(4765):747–753.

2. Jain D, Salunke DM: Antibody specificity and promiscuity. Biochem J 2019, 476(3):433–447.

3. Dimitrov JD, Planchais C, Roumenina LT, Vassilev TL, Kaveri SV, Lacroix-Desmazes S: Antibody polyreactivity in health and disease: statu variabilis. J Immunol 2013, 191(3):993–999.

4. Jaiswal D, Verma S, Nair DT, Salunke DM: Antibody multispecificity: A necessary evil? Mol Immunol 2022, 152:153–161.

5. Vashisht S, Verma S, Salunke DM: Cross-clade antibody reactivity may attenuate the ability of influenza virus to evade the immune response. Mol Immunol 2019, 114:149–161.

6. Lythgoe KA, Fraser C: New insights into the evolutionary rate of HIV-1 at the within-host and epidemiological levels. Proc Biol Sci 2012, 279(1741):3367–3375.

7. Lemey P, Rambaut A, Pybus OG: HIV evolutionary dynamics within and among hosts. AIDS Rev 2006, 8(3):125–140.

8. Andrews SM, Rowland-Jones S: Recent advances in understanding HIV evolution. F1000Res 2017, 6:597.

9. Kwong PD, Mascola JR: HIV-1 Vaccines Based on Antibody Identification, B Cell Ontogeny, and Epitope Structure. Immunity 2018, 48(5):855–871.

10. Kepler TB, Liao HX, Alam SM, Bhaskarabhatla R, Zhang R, Yandava C, Stewart S, Anasti K, Kelsoe G, Parks R et al: Immunoglobulin gene insertions and deletions in the affinity maturation of HIV-1 broadly reactive neutralizing antibodies. Cell Host Microbe 2014, 16(3):304–313.

11. Ofek G, Tang M, Sambor A, Katinger H, Mascola JR, Wyatt R, Kwong PD: Structure and mechanistic analysis of the anti-human immunodeficiency virus type 1 antibody 2F5 in complex with its gp41 epitope. J Virol 2004, 78(19):10724–10737.

12. Kwong PD, Mascola JR, Nabel GJ: Rational design of vaccines to elicit broadly neutralizing antibodies to HIV-1. Cold Spring Harb Perspect Med 2011, 1(1):a007278.

13. Corti D, Lanzavecchia A: Broadly neutralizing antiviral antibodies. Annu Rev Immunol 2013, 31:705–742.

14. Gach JS, Leaman DP, Zwick MB: Targeting HIV-1 gp41 in close proximity to the membrane using antibody and other molecules. Curr Top Med Chem 2011, 11(24):2997–3021.

15. Barras JP: [Relation of blood viscosity and hematocrit I]. Helv Physiol Pharmacol Acta 1965, 23(1):16–25.

16. Cleveland SM, Buratti E, Jones TD, North P, Baralle F, McLain L, McInerney T, Durrani Z, Dimmock NJ: Immunogenic and antigenic dominance of a nonneutralizing epitope over a highly conserved neutralizing epitope in the gp41 envelope glycoprotein of human immunodeficiency virus type 1: its deletion leads to a strong neutralizing response. Virology 2000, 266(1):66–78.

17. Reading SA, Heap CJ, Dimmock NJ: A novel monoclonal antibody specific to the C-terminal tail of the gp41 envelope transmembrane protein of human immunodeficiency virus type 1 that preferentially neutralizes virus after it has attached to the target cell and inhibits the production of infectious progeny. Virology 2003, 315(2):362–372.

18. Heap CJ, Reading SA, Dimmock NJ: An antibody specific for the C-terminal tail of the gp41 transmembrane protein of human immunodeficiency virus type 1 mediates post-attachment neutralization, probably through inhibition of virus-cell fusion. J Gen Virol 2005, 86(Pt 5):1499–1507.

19. Cheung L, McLain L, Hollier MJ, Reading SA, Dimmock NJ: Part of the C-terminal tail of the envelope gp41 transmembrane glycoprotein of human immunodeficiency virus type 1 is exposed on the surface of infected cells and is involved in virus-mediated cell fusion. J Gen Virol 2005, 86(Pt 1):131–138.

20. Mouquet H: Antibody B cell responses in HIV-1 infection. Trends Immunol 2014, 35(11):549–561.

21. Benjelloun F, Lawrence P, Verrier B, Genin C, Paul S: Role of human immunodeficiency virus type 1 envelope structure in the induction of broadly neutralizing antibodies. J Virol 2012, 86(24):13152–13163.

22. Mouquet H, Nussenzweig MC: Polyreactive antibodies in adaptive immune responses to viruses. Cell Mol Life Sci 2012, 69(9):1435–1445.

23. Chen B: Molecular Mechanism of HIV-1 Entry. Trends Microbiol 2019, 27(10):878–891.

24. Schibli DJ, Weissenhorn W: Class I and class II viral fusion protein structures reveal similar principles in membrane fusion. Mol Membr Biol 2004, 21(6):361–371.

25. Weissenhorn W, Hinz A, Gaudin Y: Virus membrane fusion. FEBS Lett 2007, 581(11):2150–2155.

26. Cowley S: The biology of HIV infection. Lepr Rev 2001, 72(2):212–220.

27. Bryson S, Julien JP, Hynes RC, Pai EF: Crystallographic definition of the epitope promiscuity of the broadly neutralizing anti-human immunodeficiency virus type 1 antibody 2F5: vaccine design implications. J Virol 2009, 83(22):11862–11875.

28. Muster T, Guinea R, Trkola A, Purtscher M, Klima A, Steindl F, Palese P, Katinger H: Cross-neutralizing activity against divergent human immunodeficiency virus type 1 isolates induced by the gp41 sequence ELDKWAS. J Virol 1994, 68(6):4031–4034.

29. Caillat C, Guilligay D, Sulbaran G, Weissenhorn W: Neutralizing Antibodies Targeting HIV-1 gp41. Viruses 2020, 12(11).

30. Barouch DH, Whitney JB, Moldt B, Klein F, Oliveira TY, Liu J, Stephenson KE, Chang HW, Shekhar K, Gupta S et al: Therapeutic efficacy of potent neutralizing HIV-1-specific monoclonal antibodies in SHIV-infected rhesus monkeys. Nature 2013, 503(7475):224–228.

31. Pantaleo G, Correia B, Fenwick C, Joo VS, Perez L: Antibodies to combat viral infections: development strategies and progress. Nat Rev Drug Discov 2022, 21(9):676–696.

32. Boughter CT, Borowska MT, Guthmiller JJ, Bendelac A, Wilson PC, Roux B, Adams EJ: Biochemical patterns of antibody polyreactivity revealed through a bioinformatics-based analysis of CDR loops. Elife 2020, 9.

33. Mouquet H, Scheid JF, Zoller MJ, Krogsgaard M, Ott RG, Shukair S, Artyomov MN, Pietzsch J, Connors M, Pereyra F et al: Polyreactivity increases the apparent affinity of anti-HIV antibodies by heteroligation. Nature 2010, 467(7315):591–595.

34. Bhowmick A, Salunke DM: Limited conformational flexibility in the paratope may be responsible for degenerate specificity of HIV epitope recognition. Int Immunol 2013, 25(2):77–90.

35. White HN, Meng QH: Diversification of specificity after maturation of the antibody response to the HIV gp41 epitope ELDKWA. PLoS One 2012, 7(2):e31555.

36. Sapkal GN, Yadav PD, Ella R, Deshpande GR, Sahay RR, Gupta N, Vadrevu KM, Abraham P, Panda S, Bhargava B: Inactivated COVID-19 vaccine BBV152/COVAXIN effectively neutralizes recently emerged B.1.1.7 variant of SARS-CoV-2. J Travel Med 2021, 28(4).

37. Gray ES, Meyers T, Gray G, Montefiori DC, Morris L: Insensitivity of paediatric HIV-1 subtype C viruses to broadly neutralising monoclonal antibodies raised against subtype B. PLoS Med 2006, 3(7):e255.

38. Perelson AS: Modelling viral and immune system dynamics. Nat Rev Immunol 2002, 2(1):28–36.

39. Sanjuan R, Domingo-Calap P: Mechanisms of viral mutation. Cell Mol Life Sci 2016, 73(23):4433–4448.

40. Jaiswal D, Kumar U, Gaur V, Salunke DM: Epitope-directed anti-SARS-CoV-2 scFv engineered against the key spike protein region could block membrane fusion. Protein Sci 2023, 32(3):e4575.

41. Halgren TA, Murphy RB, Friesner RA, Beard HS, Frye LL, Pollard WT, Banks JL: Glide: a new approach for rapid, accurate docking and scoring. 2. Enrichment factors in database screening. J Med Chem 2004, 47(7):1750–1759.

42. Roos K, Wu C, Damm W, Reboul M, Stevenson JM, Lu C, Dahlgren MK, Mondal S, Chen W, Wang L et al: OPLS3e: Extending Force Field Coverage for Drug-Like Small Molecules. J Chem Theory Comput 2019, 15(3):1863–1874.

